# Dynamical A*β*-Tau-Neurodegeneration Model Predicts Alzheimer’s Disease Mechanisms and Biomarker Progression

**DOI:** 10.64898/2026.01.27.701320

**Authors:** Pavanjit Chaggar, Jacob W. Vogel, Travis B. Thompson, Roxana Aldea, Olof Strandberg, Erik Stomrud, Sebastian Palqmvist, Rik Ossenkoppele, Saad Jbabdi, Stefano Magon, Gregory Klein, Alzheimer’s Disease Neuroimaging Initiative, Niklas Mattson-Carlgren, Oskar Hansson, Alain Goriely

## Abstract

Alzheimer’s disease is characterised by the pathological interaction of two proteins, amyloid-beta (Aβ) and tau, which collectively drive neurodegeneration and cognitive decline. The progression of Aβ, tau, and neurodegeneration biomarkers is captured by the ATN framework, which is a powerful tool for disease classification. However, since the ATN framework is mainly descriptive, it cannot quantify or predict relationships between biomarkers over time. We address this limitation by introducing a dynamical ATN (dATN) model that mechanistically simulates the spatiotemporal progression of Aβ, tau, and neurodegeneration. The dATN model integrates mechanisms of prion-like protein aggregation of Aβ and tau, network-based tau propagation, Aβ-driven catalysis of tau progression, and tau-driven neurodegeneration. We calibrated the model using multimodal longitudinal imaging data from both the ADNI and BioFINDER-2 cohorts and show that it accurately fits longitudinal regional Aβ, tau, and neurodegeneration data. Using the dATN model, we show that Aβ-induced effects predict Braak-like cortical tau progression, that the spatial colocalisation of Aβ and tau is a crucial biomarker of disease acceleration, and that tau-driven atrophy strongly correlates with observed neurodegeneration. Furthermore, by integrating the disease progression model with pharmacokinetic–pharmacodynamic simulations, we present a powerful tool that facilitates regional evaluation of therapeutic strategies targeting Aβ, identification of critical intervention windows, and prediction of heterogeneous treatment effects across brain regions. This framework unifies mechanistic understanding with clinical imaging biomarkers, offering a quantitative approach for forecasting disease progression, testing mechanistic hypotheses, and optimising personalised treatment strategies in AD.

**One Sentence Summary:** Colocalisation of A*β* and tau predicts regional tau progression and optimal A*β*-targeted intervention windows, while tau predicts neurodegeneration.

## INTRODUCTION

Alzheimer’s Disease (AD) is a multifaceted disease characterised by several interacting pathological processes which culminate in dementia. The two principal drivers of AD pathology are toxic forms of A*β* and tau proteins, which aggregate and spread throughout the brain, resulting in brain neurodegeneration and cognitive dysfunction (*1, 2*). Imaging biomarkers for A*β* (*3, 4*) and tau (*5*–*9*) are available using positron emission tomography (PET), which allow for regional quantification of pathological protein load throughout the brain and personalised AD staging. Meanwhile, structural magnetic resonance imaging (sMRI) can be used to measure neurodegeneration (*10*–*12*). The use of imaging biomarkers for quantitatively diagnosing and staging AD manifested first in the ATN framework (*13*), describing the progression of A*β*, tau and neurodegeneration, and more recently in the revised diagnosis and staging framework (*14*). However, the frameworks provide only a phenomenological description of biomarker progression and do not capture the mechanistic relationships between biomarkers. Therefore, an unmet need remains in developing a model of AD biomarker progression that provides deeper insight into disease mechanisms and enables personalised predictions of biomarker progression.

Several key features of underlying pathology have been uncovered through animal models and human observational studies that may inform the progression of AD biomarkers from underlying neurobiology (*15*). First, both A*β* and tau exhibit distinct and complementary spatial profiles. A*β* is diffusely present throughout the cortex before spreading to subcortical regions, such as the medial temporal lobe (*16, 17*), with temporal differences in accumulation largely resulting from spatial heterogeneity in maximal A*β* load (*18*). In contrast, tau undergoes more systematic staging, with cross-sectional studies showing that pathology begins first in the entorhinal cortices, then spreads sequentially throughout the temporal lobe, inferior parietal and posterior cingulate structures (*19, 20*). Second, mounting evidence from animal models (*21*–*24*) and modelling of human neuroimaging data (*25*–*27*) supports the role of network-based transsynaptic spread in the sequential progression of tau pathology through the brain. Third, while the initial accumulation of tau in the medial temporal lobe is common during the ageing process, characterised as primary age-related tauopathy (PART) (*28, 29*), animal studies show that the presence of A*β* results in faster tau tangle formation, accumulation and spread (*30*–*33*). This is also supported by in-vivo human studies, showing that tau spread and aggregation is accelerated by A*β* deposition (*34, 35*) and that *A*^+^*T*^+^ individuals are at greater risk of cognitive decline compared to *A*^+^*T*^−^ and *A*^−^*T*^−^ individuals (*36*). Fourth, neuroimaging studies have shown that neurodegeneration is more strongly correlated with tau burden than with A*β* burden (*37*–*39*), and that both neurodegeneration and tau are correlated with cognitive decline (*40*–*42*). As yet, there are limited tools available for tying together these putative mechanisms of AD pathology to the progression of ATN biomarkers in humans.

Previous modelling work has focused either on single biomarkers, such as A*β* or tau (*18, 26, 27*), pairwise combinations of ATN biomarkers (*43, 44*), or summary values of ATN biomarkers (*45*), thereby either neglecting potentially important interaction mechanisms or losing the spatial information available from neuroimaging. Operationalising the complete spatiotemporal ATN pathway will facilitate hypothesis testing from biomarker data, and provide better characterisations and predictions of biomarker progression. Modelling and simulation may also help address challenges present in the burgeoening field of personalised AD intervention. Current disease modifying treatments for AD work by eliminated A*β* plaques from the brain (*46, 47*) and show modest but significant effects on disease slowing. However, the mechanisms by which A*β* removal affects tau and neurodegeneration, and if or how these effects may contribute to disease slowing, remain unknown. Recent work has sought to combine pharmacokinetic and pharmacodynamic models of A*β* progression to analyse clinical trials (*45, 48, 49*). However, the models include limited information from biomarkers, modelling only A*β* centiloid (CL) (*50*) evolution and therefore excluding regional information, as well as information available from tau PET and sMRI scans. Given the complex spatiotemporal relationships among A*β*, tau, and neurodegeneration, it will be important to fully model temporal and spatial information on biomarker development to better assess and monitor changes in pathology in response to intervention.

Here, we present a modelling and simulation framework that can be used to consolidate existing knowledge concisely, gain further mechanistic insight from biomarker data, and offer translational benefits through integration with intervention therapies. We build on previous work (*27, 43*) to develop the dynamical ATN (dATN) model, which integrates our knowledge about A*β*, tau and neurodegeneration interactions using dynamical systems. We validate the model using multimodal longitudinal imaging biomarkers from the Alzheimer’s Disease Neuroimaging Initiative (ADNI) and the BioFINDER-2 (BF2) studies, demonstrating its ability to capture AD-specific progression of ATN biomarkers. Simulations with the dATN model offer mechanistic insight by showing that A*β* promotes the cortical Braak staging of tau spread via local interactions and that the acceleration of AD progression follows the spatial colocalisation of A*β* and tau. The dATN model architecture facilitates integration with other biomarkers and domains. Here we integrate the dATN model with pharmacokinetics/pharmacodynamics (PK/PD) to provide a spatiotemporal model of the regional downstream effects of A*β* targeting therapies on the ATN cascade. We demonstrate the translational utility of the dATN model by showing how modelling longitudinal ATN biomarkers can help to optimise the effects of anti-A*β* therapies. Overall, the dATN model provides the necessary quantitative and dynamical framework to consolidate existing knowledge, facilitate hypothesis testing with complex spatiotemporal data, and enable mechanism-based biomarker forecasting and treatment optimisation in AD.

## RESULTS

### Dynamical ATN Model Predicts that Tau Staging Follows A*β* Heterogeneity

The dATN model rests on the following physical assumptions based our current understanding of the ATN disease process: (i) A*β* and tau follow a regionally heterogeneous prion-like autocatalytic process of protein accumulation; (ii) A*β* does not undergo a significant transport process (measurable through neuroimaging); (iii) Tau transport takes place along axonal connections; (iv) A*β* accelerates tau accumulation; (v) Tau deposition results in structural neurodegeneration.

The model predicts that the regional tau load is determined by a combination of A*β*-independent and A*β*-dependent factors (although the tau levels resulting from A*β*-independent effects are likely to be minimal, and are assumed to be PART). We test this prediction in multiple ways. First, in Fig. 1A and Fig. 1B we show the SUVR carrying capacities for florbetaben (FBB) A*β* PET and flortaucipir (FTP) PET and in Fig. 1C, we show that there is a linear correlation between these values for a combination of A*β* and tau tracers used in ADNI and BF2. Second, we use the linear coefficient between FBB and FTP to simulate tau concentration in each Braak region using the dATN model, and in Fig. 1D, we show that the dATN model predicts that regional tau progression driven by regional A*β* load and network-based tau spread follows expected Braak staging. Lastly, in Fig. 1E, we show the regional trajectories of A*β* SUVR, tau PET SUVR and neurodegeneration. The model captures the expected sequential progression of A*β*, tau and neurodegeneration. More importantly, the regional trajectories of tau reflect those observed using tau PET, namely, early accumulation in the lateral temporal, medial parietal, lateral parietal and lateral frontal cortices. Furthermore, when compared to the trajectories shown in Fig. S1, in which there is no catalytic effect of A*β* on tau, we see that the model predicts that A*β* is necessary for cortical progression of tau. These results, derived from the dATN model and cross-sectional data, suggest that Braak-like staging and cortical heterogeneity of tau emerge from the heterogeneity in tau acceleration given by regional A*β* load, indicating that A*β* orchestrates the cortical progression of tau across the brain network. This initial analysis demonstrates how the dATN enables the extraction of testable hypotheses and parameters related to biomarker evolution. We next calibrate the model using individual-level longitudinal data to further test its predictions.

**Fig. 1.**
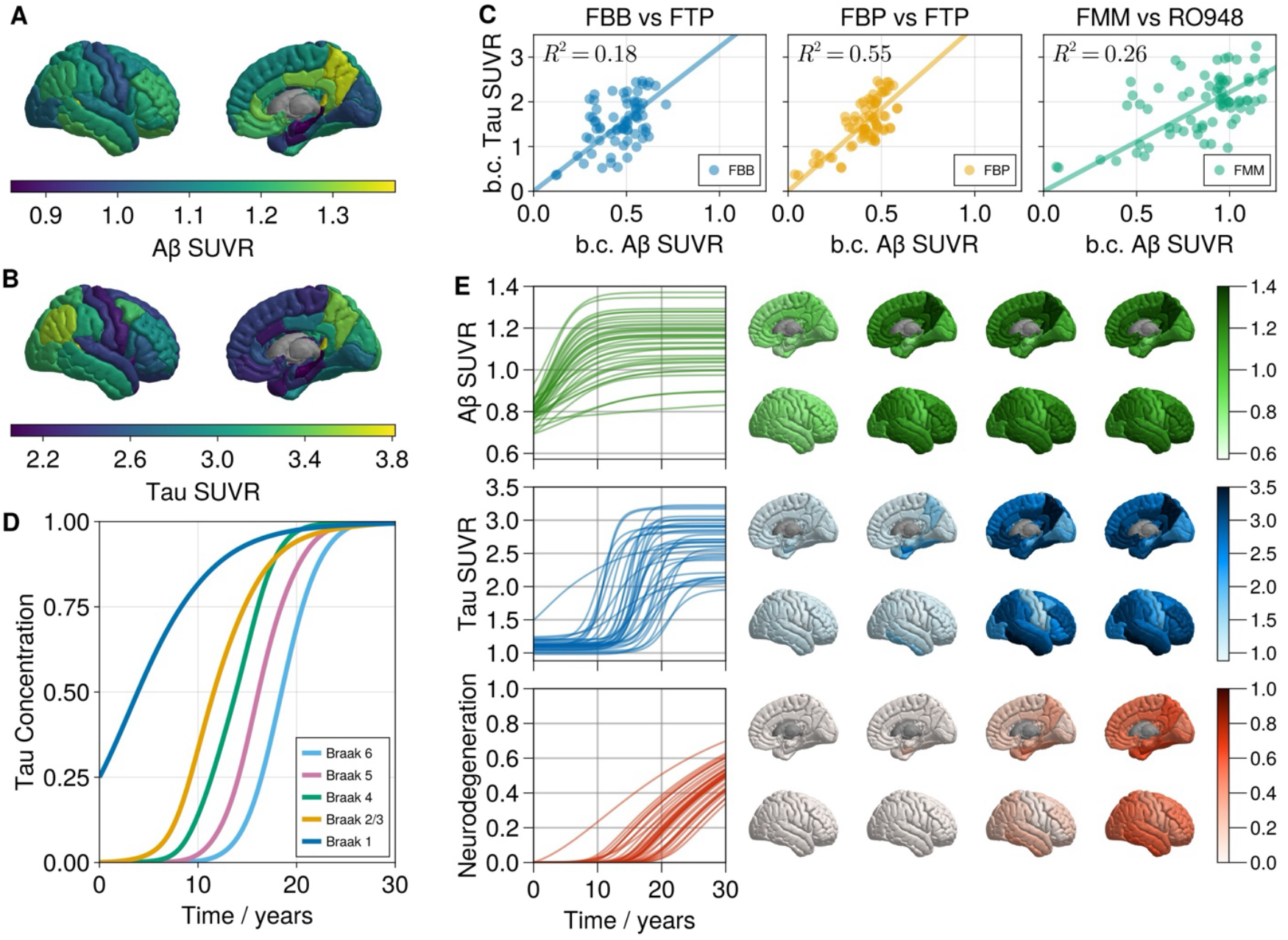
Dynamics of the ATN model. (A) Estimated carrying capacities for Aβ measured by florbetaben (FBB) PET on the right hemisphere. (B) Estimated regional carrying capacities for tau measured by flortaucipir (FTP) PET on the right hemisphere. (C) Relationship between baseline corrected Aβ PET SUVR (carrying capacity SUVR minus baseline SUVR) for FBB, florbetapir (FBP) and flutemetamol (FMM) compared to inferred baseline corrected tau PET SUVR carrying capacities (carrying capacity SUVR minus baseline and PART SUVR) for FTP and RO948. (D) Summary of simulation in (E) for tau concentration over the Braak regions. Tau concentration is calculated by scaling the SUVR between the regional baseline values and carrying capacities. (E) Simulation from the dATN model (Equation 12) using seeding of Aβ throughout the cortex of A_i_(0) = 0.2)A_∞,i_ − A_0,i_+ for i = 1, …, R, where A_∞,i_ and A_0,i_ are regional carrying capacities and baseline values, respectively, for FBB, and a tau seeding of T_i_(0) = 0.2)T_∞,i_ − T_0,i_+ in the bilateral entorhinal cortex, i = {27,62} and T_i_(0) = T_0,i_ elsewhere, where T_∞,i_ and T_0,i_ are regional carrying capacities and baseline values, respectively, for FTP. We use synthetic parameters, with tau in a production-dominated regime, α = 0.75/yr, ρ = 0.025/yr, γ = 0.5/yr, β = 3.21, η = 0.05/yr. The trajectories for the right cortex are shown over a 30-year period, and cortical renderings on the right cortex are shown at t = {0,10,20,30}.

### Forecasting Multi-modal Longitudinal Biomarkers

We use longitudinal A*β* PET, tau PET and sMRI data available in ADNI and BF2 to test whether the dATN model accurately describes biomarker progression. We use *A*^+^*T*^+^ subjects who have at least two A*β* PET scans and at least three tau PET scans. In total, there are *N* = 34 subjects in ADNI and *N* = 48 subjects in BF2, with mean duration between baseline and final scan of 3.6 and 3.3 years for A*β* and tau PET, respectively, in ADNI, and 4.2 years for A*β* and tau PET in BF2. We use a hierarchical probabilistic model with population-level and individual-level rate parameters and a population-level A*β*/tau coupling parameter (see Methods for the detailed model specification). The posterior distributions of population-level parameters are shown in Fig. 2A and the individual-level posterior distributions are shown in Fig. S2. All parameters are well identified and there is good agreement in the model parameters between the ADNI and BF2 cohorts, suggesting that the mechanisms in the model are generalisable across the different cohorts and different tracers. Interestingly, the parameter distributions show that the transport and production parameters for tau, *ρ* and *γ*, are of a similar magnitude and not dominated by a single process. However, tau progression can be considered production-dominated (*ρ* ≪ *γ*) when accounting for the catalytic effect of A*β*, implying that A*β* accelerates tau production and tau-induced neurodegeneration.

**Fig. 2.**
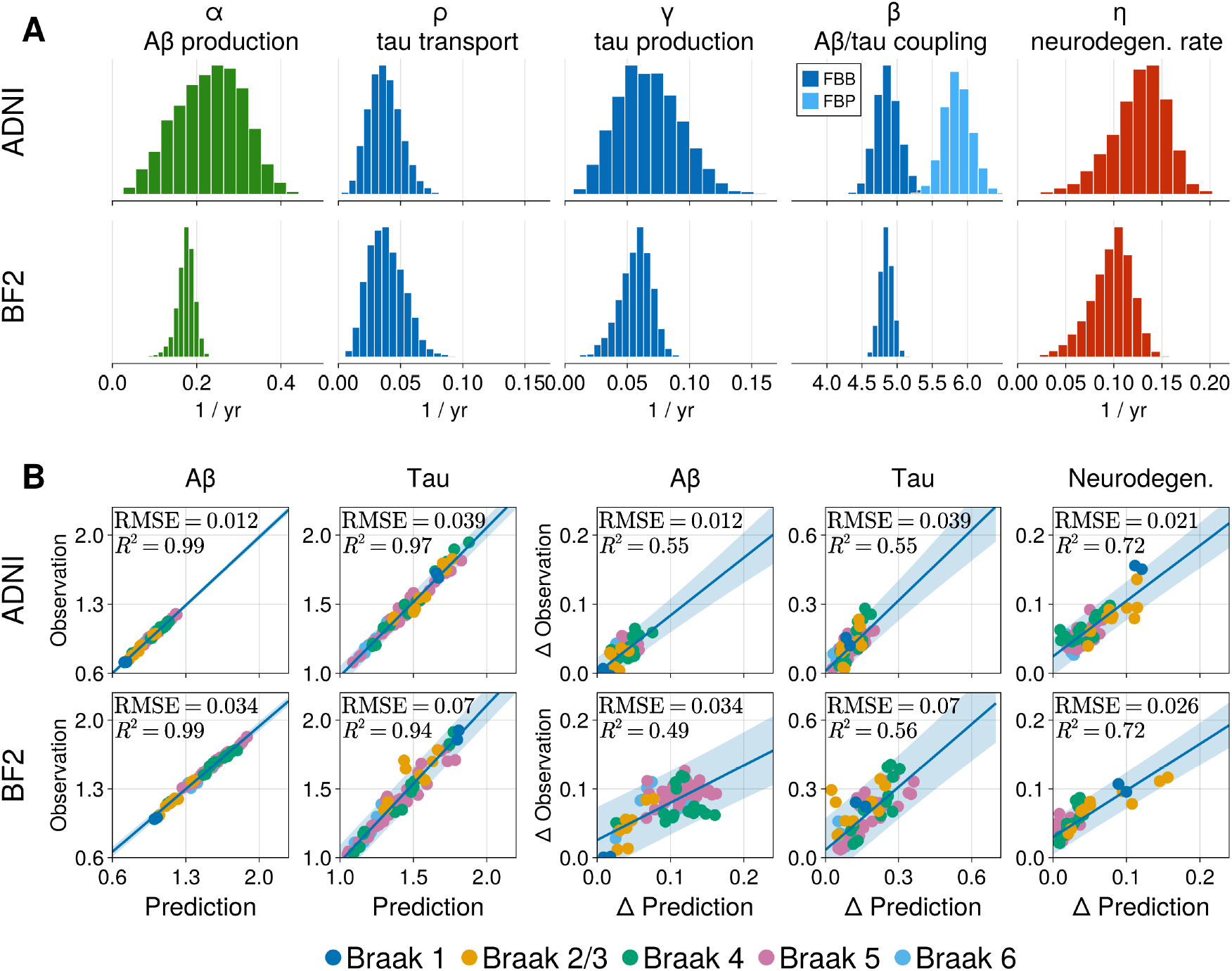
Posterior predictions from the ATN model calibrated to A^+^T^+^ subjects in ADNI and BF2. (A) Posterior distribution for dATN model population-level parameters from the ADNI cohort (top) and the BF2 cohort (bottom). (B) Posterior model fit for ADNI (top) and BF2 (bottom), averaged over subjects per regions, using personalised posterior parameters (Fig. S2). The first two panels show the predicted vs observed SUVR for Aβ and tau for the final longitudinal scan. The remaining three panels show the predicted vs observed change in Aβ SUVR (left), tau SUVR (right), and neurodegeneration between baseline and final scan. Neurodegeneration is measured as change in volume relative to baseline scan (normalised by total intracranial volume and baseline scan), therefore, the observed vs predicted is the same as the observed change vs predicted change, and the former is therefore excluded. Each point represents a region in the DK parcellation averaged over subjects. Regions are colored by their Braak stage designation.

The model fit averaged over subjects for each DK atlas region is shown in Fig. 2B. For all ATN biomarkers, the model provides a good fit to data, with *R*^2^ values for predictions vs observations of 0.99, 0.97 for A*β* and tau, respectively, in ADNI, and 0.99, 0.94, respectively, in BF2. The model also performs well at predicting longitudinal changes from baseline biomarkers. For A*β* and tau and neurodegeneration, the *R*^2^ for the predicted vs observed longitudinal change are 0.55, 0.55, and 0.72 respectively, in ADNI, and 0.49, 0.56 and 0.72, respectively, for BF2. These results show that regional A*β* dynamics predict heterogeneity in regional tau burden and change. Furthermore, we find that change in neurodegeneration from baseline is strongly correlated with regional tau dynamics throughout Braak stage regions, demonstrating a mechanistic link between A*β*-induced tau dynamics and neurodegeneration. The model fits the ADNI data more accurately than the BF2 data, likely due to demographic differences in the cohorts, with individuals in BF2 being dominated by MCI subjects with a lower average CL (mean ± SD = 73 ± 32), compared to ADNI which is dominated by AD subjects with a higher average CL (mean ± SD = 84 ± 34). Furthermore, the PET tracers used in BF2 have a higher dynamic range than those used in ADNI and, therefore, a greater absolute change in SUVR may not represent greater change in disease. Since subjects in BF2 cohort are, on average, earlier in the disease process than those in the ADNI cohort, the observed change in ATN biomarkers may not wholly represent AD pathology and are therefore not as well captured by the dATN model. Overall, the dATN model, based on physical assumptions about ATN pathologies and using pooled coupling between A*β* and tau accurately fits regional longitudinal ATN biomarker data, validating the model’s predictions that regional A*β* drives cortical tau progression and demonstrating that tau results in neurodegeneration.

### A*β*/Tau Colocalisation Separates Lag-phase and Acceleration-phase of Tau

We next use the calibrated dATN model to investigate the spatiotemporal nature of A*β*/tau interactions, specifically *when* and *where* in the brain they colocalise such that A*β* induces a significant accelerating effect on tau. To do so, we simulate from the dATN model (Equation 14) using the inferred mean individual-level parameters with initial conditions from an early tau group in ADNI, defined by a mean CL ≥ 40, when A*β* is progressing and initial tau deposition is observable through PET (*51, 52*).

To quantify the colocalisation of A*β* and tau, we develop a derivative-based method for determining A*β* and tau colocalisation thresholds: for A*β* we chose the regional concentration at which the regional velocity is maximised (when A*β* load is changing the fastest and exerting a large and increasing effect on tau); for tau we choose the regional concentration at which the regional jerk is maximised (when acceleration is changing the fastest). These are shown for A*β* and tau in Fig. 3A and Fig. 3B, respectively, for the left inferior temporal lobe (IT), along with the simulated trajectories in the left hemisphere regions of the DK atlas, with the IT and entorhinal cortex (EC) highlighted. In the left hemisphere, the inferior temporal lobe is the first to meet the detection threshold for both A*β* and tau, despite having similar levels of initial A*β* and significantly less tau at baseline compared to the EC. Furthermore, the progression of tau in cortical regions follows the colocalisation point, suggesting that A*β*/tau colocalisation precipitates widespread tau progression. In Fig. 3C we show the full colocalisation order, the temporal sequence in which regions reach their A*β* and tau thresholds, for the left and right hemispheres using data from ADNI and BF2. In the left hemisphere, the initial colocalisation occurs in IT for ADNI and BF2. In the right hemisphere, initial colocalisation occurs in the banks of the superior temporal sulcus for ADNI, and in the IT for BF2. Subsequent cortical tau accumulation after initial colocalisation is seen predominantly in cortical regions of high amyloid deposition, namely the lateral temporal lobes, precuneus, inferior parietal and lateral frontal cortices (see Table S1 and S2 for the full colocalisation order in ADNI and BF2, respectively). In Fig. 3D, we show the probability for a given site to be the first region of A*β*/tau colocalisation for ADNI and BF in the left and right hemisphere, which shows slight variations in initial colocalisation throughout the temporal cortices, but is dominated by the mean inferred colocalisation sites. This suggests that slight deviations in dATN parameters are unlikely to change the initial colocalisation site. The full list of regions and their colocalisation probabilities is provided in Tables S3 and S4, for ADNI and BF2, respectively. Through this analysis, we identify the lateral temporal lobes as the site of initial colocalisation and, except for the right hemisphere in ADNI, the IT is the most likely region for initial colocalisation. Overall, these results highlight that the locus of A*β*-induced tau acceleration likely resides in the lateral temporal lobe and immediately precedes the progression of tau throughout the cortex. The colocalisation point, therefore, demarcates a window of slow tau accumulation following tau seeding, after which there is a cascade of cortical tau acceleration.

**Fig. 3.**
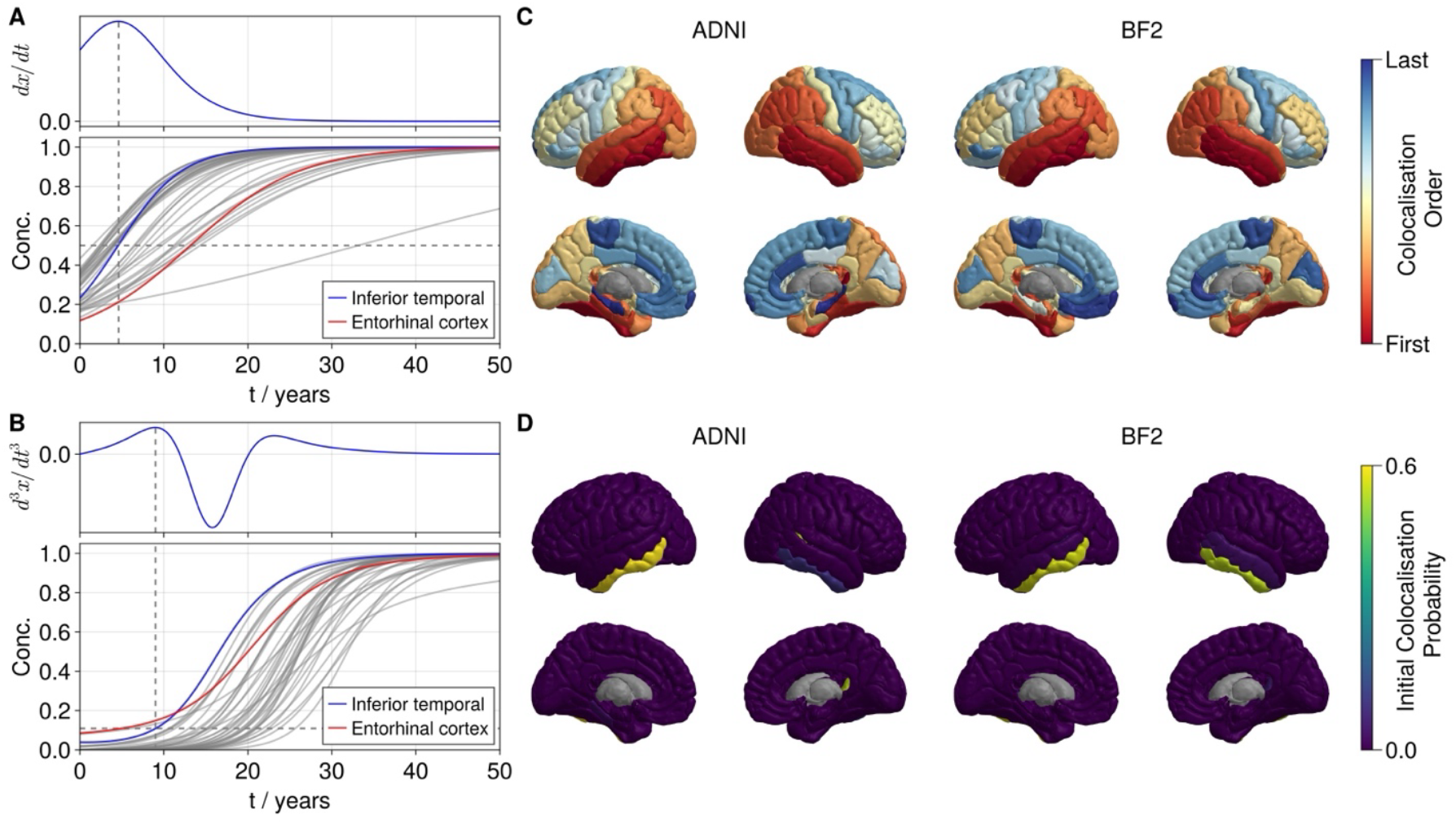
Quantifying Aβ/tau colocalisation. (A) Simulated trajectory of Aβ concentration for regions in the left hemisphere of the DK atlas (bottom). The IT is highlighted in blue and the EC is highlighted in red. The detection threshold for a region is determined as the point of maximum velocity, shown in the top panel for the IT. (B) Simulated trajectory of tau concentration for regions in the left hemisphere of the DK atlas (bottom). The IT is highlighted in blue and the EC is highlighted in red. The detection threshold for a region is determined as the point of maximum jerk during the accumulation phase (top). (C) Order of Aβ/tau colocalisation using mean posterior model parameters calculated separately on the left and right hemisphere for ADNI and BF2. Warm colors represent early colocalisation and cool colors represent late colocalisation. (D) Probability of a region being the initial site of Aβ/tau colocalisation calculated separately using on the left and right hemisphere for ADNI and BF2. In the left hemisphere, the initial colocalisation occurs most often in IT for ADNI and BF2. In the right hemisphere, initial colocalisation occurs most often in banks of the superior temporal sulcus for ADNI and in the IT for BF2.

### Colocalisation predicts optimal intervention strategies in-silico

Given the ability of the dATN model to predict longitudinal biomarker changes at the regional level, it may provide utility in simulating ATN biomarkers in response to intervention. Therefore, we next investigate the potential for the ATN model to inform therapeutic design with A*β* targeting therapies by coupling the calibrated dATN model with a synthetic PKPD model of drug entry and action in the brain. We assume that drug entry to the brain is dominated by distance-based diffusion from major blood vessels surrounding the choroid plexus (*53*). Furthermore, we assume that upon A*β* removal, the carrying capacity of tau remains at the highest level prior to A*β* removal. A simulation from the PKPD model in the left hemisphere is shown in Fig. 4A, in which a dose of the drug is administered monthly for a 30-year period. Under the administration constraints, the drug accumulates to a high concentration in regions close to the choroid plexus and saturates at a much lower rate in regions further away. We simulate from the coupled dATN-PKPD model in the left hemisphere using mean individual-level posterior parameters and initial conditions derived from the early tau group, shown in Fig. 4B. In Fig. 4C, we simulate from the coupled dATN-PKPD model to examine how A*β* targeting drugs affect the trajectories of A*β*, tau and neurodegeneration, showing both the placebo case (left panel) and active case (right panel). In the top right panel of Fig. 4C, we show the effect of A*β* targeting treatment on A*β* concentration, and we see that there is a sharp decline of A*β* in regions proximal to the choroid plexus, where drug concentration is high, and a slower decline in regions distal to the choroid plexus. As a result, there is heterogeneous clearance of A*β*, with the drug efficacy decaying with distance from the entry locations. Downstream, there is a slowing of tau progression and neurodegeneration. However, the model predicts suboptimal tau elimination in the temporal lobes, due to age-related effects (PART) and persistent A*β* effects which are assumed to be present but not accumulating after A*β* elimination. To further examine how intervention with A*β* targeting therapies affects ATN biomarker progression, we simulate intervention at different points across the AD timeline with the same parameters and initial conditions. In Fig. 4D we show the simulated trajectories of the dATN-PKPD model averaged over Braak stages 1-3, with intervention starting at different time points along the AD continuum (*t*_0_ = 0,5,10,20 years) and continuing onwards with doses administered every month until *t* = 30 years. For all interventions, A*β* is completely removed within two years of treatment onset. For intervention starting at *t*_0_ = 0 there is a 75% reduction in tau concentration and 50% reduction in neurodegeneration at 30 years compared to placebo. With delayed intervention, the dATN-PKPD model predicts non-linear increases in tau and neurodegeneration at 30 years, with late intervention (*t*_0_ = 20) offering a negligible reduction compared to placebo. This effect is emphasised in Fig. 4E, which shows the endpoints for tau and neurodegeneration following simulated treatment with different initial dosing times, starting at successive 2-year intervals from *t*_0_ = 0, and indicates a non-linear increase in tau and neurodegeneration that saturates towards the placebo case. The relative change in endpoint tau and neurodegeneration between successive initial administration times is shown in Fig. 4F. We can see that there are accelerating increases in tau and neurodegeneration as treatment is delayed before an inflection, after which relative changes in end state tau and neurodegeneration decays exponentially toward placebo. Strikingly, the inflection point for relative change in tau and neurodegeneration coincides with the colocalisation of A*β* and tau (highlighted by the solid vertical line). The dATN-PKPD model therefore predicts that after there are diminishing returns from intervention with A*β*- targeting therapies after colocalisation occurs. To test this result against sensitivity to initial conditions, we rerun this analysis with initial conditions from groups earlier or later in the AD continuum, shown in Fig. S3, which indicates that the inflection point in outcomes coincides with colocalisation when cortical tau is present (≥ 40 CL). Therefore, A*β*/tau colocalisation may represent a critical window for interventional success, where there is enhanced outcome benefit for treatment pre-colocalisation, and diminishing outcome benefit for intervention post-colocalisation, suggesting early intervention is especially important when considered over time horizons consistent with the length of AD progression.

**Fig. 4.**
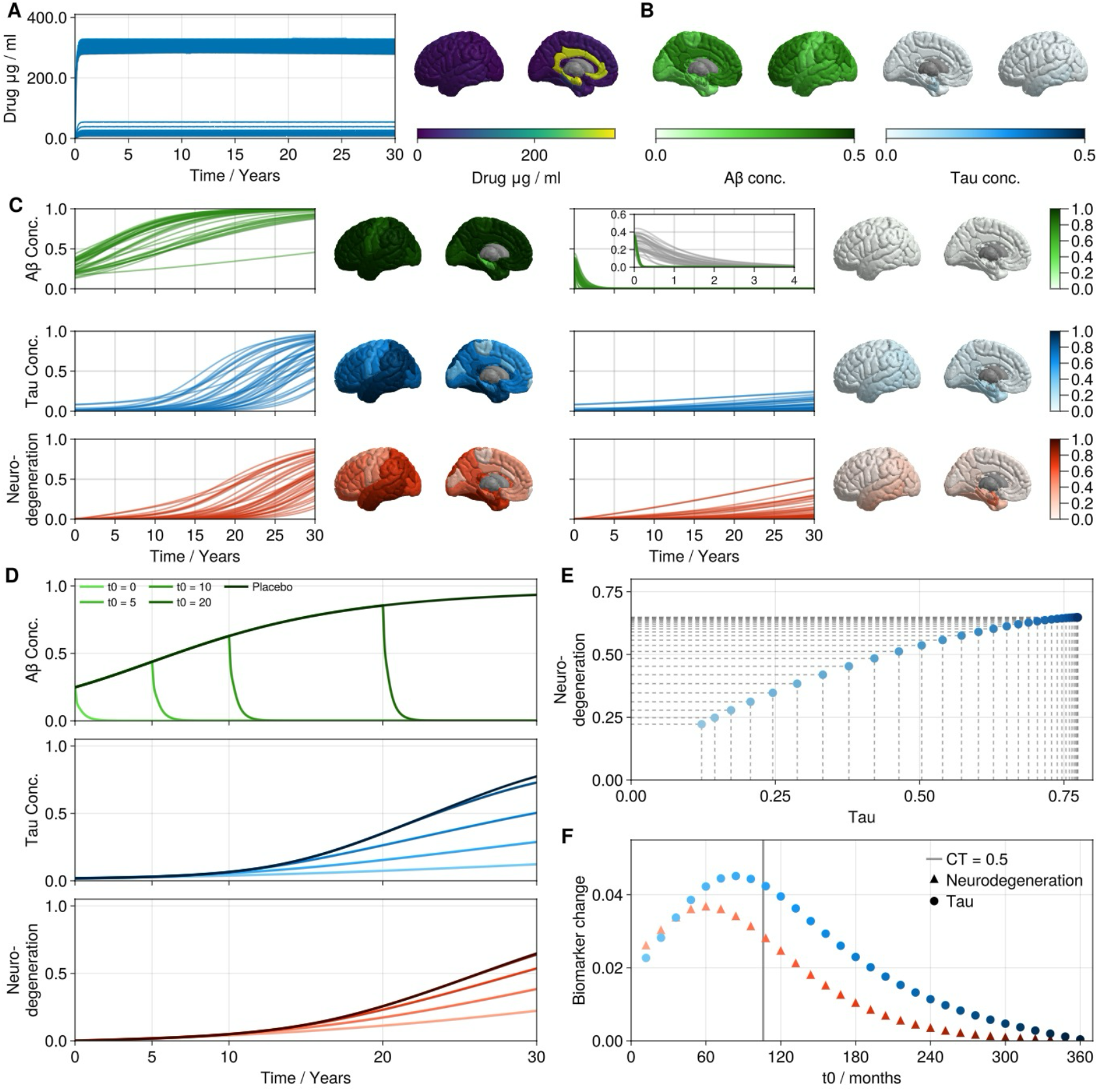
Simulated effects of Aβ targeting therapy on ATN biomarkers. (A) Simulated progression of drug concentration in the brain, assuming diffusion from dominant blood vessels surrounding lateral ventricles (54). (B) Average FBB (left) and FTP (right) concentrations from the early tau group (defined by a mean CL ≥ 40, see methods) used as initial conditions for the dATN-PKPD model (C) Simulated progression of Aβ, tau and neurodegeneration under placebo conditions (left) and with active drug effects (right) from the dATN-PKPD model (Equation 28). Both simulations start with the initial conditions in (B) and zero baseline neurodegeneration, with mean individual-level parameters from the calibrated dATN model. All parameters are provided in Table 3. (D) Simulated average trajectories of ATN biomarkers for a Braak stage 1-3 composite region with different initial treatment onset times, using parameters in Table 3. (E) Simulated neurodegeneration and tau levels at t=30 years months for interventions starting at different time points, with lighter colors representing earlier intervention. (F) simulated relative increase in neurodegeneration and tau between different initial treatment onset times.

We have shown that a simple model of drug administration, uptake and interaction provides insight into the spatiotemporal dynamics of A*β*-targeting therapies. Furthermore, when paired with the dATN model, we allow for a spatial examination of AD intervention that may provide a useful tool for clinical and pharmaceutical scientists in designing effective administration protocols.

## DISCUSSION

Substantial research has pointed towards mechanistic links between A*β*, tau and neurodegeneration, culminating in the A*β* cascade hypothesis and the ATN framework, which qualitatively summarise our knowledge of ATN interactions and their biomarker progressions. Here, we have formalised this knowledge into a compact mathematical model that clearly distils our current knowledge about ATN interactions. In so doing, we show that the spatiotemporal complexity of AD can emerge from only a few ingredients, namely regional A*β* heterogeneity, axonal tau spread, A*β*-induced tau acceleration, and tau induced neurodegeneration. From these ingredients, we show that the model reproduces the Braak-like dynamics expected in tau staging and can accurately fits longitudinal ATN biomarker data. We further demonstrate that the model can provide mechanistic insight into disease processes through the identification and quantification of A*β*-tau colocalisation, which may have translational benefits through optimising intervention strategies with A*β* modifying treatments. Overall, this work presents a quantitative framework for describing, simulating and interrogating AD pathology through ATN biomarker progression.

We have previously demonstrated the effectiveness of a regionally specific model of tau production and spread and highlight the importance of regionally varying parameters in accurately simulating the progression of tau pathology (*27*). We proposed that A*β* is the most likely candidate for providing a mechanism for heterogeneous tau dynamics since A*β* itself has distinct spatial topography, is correlated with neocortical tau pathology (*20*), and is known to enhance tau pathology (*30*–*33*). Here, we have shown that A*β* can promote regional progression and accumulation of tau that accurately fit longitudinal tau PET and neurodegeneration progression, supporting evidence for the mechanistic links between A*β* and tau as co-conspirators in promoting AD pathology and neurodegeneration. Interestingly, our results show that a population-averaged A*β*-tau coupling parameter is sufficient to accurately predict longitudinal ATN biomarker progression, suggesting that heterogeneity in A*β*/tau interaction strength may be limited. However, the value of the coupling parameter is higher than we might expect it to be. Through linear regression, we estimated relationship between A*β* carrying capacities and tau carrying capacities to be a scale factor of between 2.3-3.6, whereas the mean value from the Bayesian analysis is between 4.8-6.0. This could reflect at least three possibilities: (1) the tau carrying capacities inferred via mixture modelling are systematically lower due to insufficient sampling of end stage disease, (2) late stage neurodegeneration results in a decreased regional maximal SUVR that is not accounted for by the dATN model, (3) the A*β* mechanism is being overleveraged to account for changes in tau, where other mechanisms might better fit. Furthermore, we note that A*β* does not entirely explain regional tau dynamics and other factors might influence regional tau. Other candidates that might act together with A*β* to influence tau, including genetic risk factors (*54, 55*), such microtubule association protein tau (*55*–*59*) expression, and gradients in functional activation that have been shown to influence tau progression (*60*–*62*). Therefore, while our results suggest a strong mechanistic link between A*β* and tau and neurodegeneration data, future work should aim to include multiple risk factors influencing tau pathology, which can be easily implemented within our framework.

A major result in the current work is being able to identify at regional level the spatiotemporal location of A*β*/tau colocalisation. The dATN model predicts the inferior temporal lobe is the most likely region for colocalisation to occur, and advances on previous work (*34*) to provide a model that can simulate multiple trajectories based on real-world parameters that fully characterises a colocalisation order, spreading from the inferior temporal gyrus to the fusiform, precuneus, inferior parietal and lateral frontal regions. Additionally, we observe asymmetry in possible colocalisation order and sites between the left and right hemispheres, which may underpin A*β* induced hemispheric asymmetry in tau burden (*63*). After initial colocalisation, the regions following the inferior temporal lobe coincide with those cortical regions particularly prone to neurodegeneration (*64*), suggesting that early A*β*/tau colocalisation may correlate with accelerated neurodegeneration. The dATN model builds on existing methods to extend and generalise the notion of A*β*/tau colocalisation to be able to predict colocalisation at an individual level and may be useful for predicting areas of neurodegeneration and subsequent cognitive impairment.

There have been several studies showing that A*β* is necessary for the acceleration of tau and the progression of tau pathology into the neocortex (*28, 30, 34, 51*), however, the precise mechanism by which this occurs are unclear. Due to the spatial segregation of A*β* and tau during AD (*16, 20*), A*β* does not have an immediate local effect on tau pathology. Recently, work has pointed toward a remote interaction of A*β* on tau. In Lee et al. 2022 (*34*), the authors propose a model in which tau propagation is catalysed by A*β* in two phases, first by remote A*β* effects on the EC that promote the *outward* spread of tau to cortical areas of high A*β*, and second by local interactions in which A*β* catalyses tau production, starting in the inferior temporal lobe. This is complemented by recent work from Roemer-Cassiano et al. 2025 (*61*), who show that tau spread into the cortex may be facilitated by A*β*-induced cortical hyperconnectivity. However, studies by Giorgio et al. 2023 (*65*) and Alexandeerson et al. 2025 (*66*) propose a different kind of remote effect in which cortical A*β* accelerates early MTL tau accumulation through hyperexcitability. The model presented here does not include any remote interactions of A*β* on tau, however, it is nonetheless able to predict AD progression and fit longitudinal data. This suggests that remote connections may not be necessary to explain the preferential invasion of tau to regions of high A*β*, and that local A*β*/tau interactions are sufficient. This is also shown by similar work using model selection methods to show that remote interaction is not necessary to predict longitudinal biomarker data (*44*). The inconclusive results regarding the relationship between A*β* and tau from biomarker data likely stems from the relatively coarse nature of PET imaging data that does not entirely represent pathology, since tracers bind to specific aggregated forms of A*β* and tau, and therefore cannot wholly capture the rich dynamics of A*β* and tau at various spatial and temporal scales, such as the accumulation of smaller oligomers of A*β* and tau, at which different interaction effects may occur (*67, 68*). Overall, while our results suggest that a local interaction between A*β* and tau is sufficient to explain the cortical spreading of tau, we are limited in drawing strong mechanistic conclusions on the nature of A*β*/tau interactions from the model and data used in this study. However, the dATN framework is readily adaptable, and future work may seek to extend the dATN model to include biomarkers of other processes, such as fluid biomarkers, FDG-PET or functional MRI, that may elucidate the effects of different processes on A*β*/tau interactions.

By coupling the dATN model with a PKPD model, we show that the time and location of A*β*/tau interactions could be crucial for the optimal application of disease-modifying interventions in AD. Current therapies, such as Lecanemab and Donanemab, focus on the reduction of A*β* (*46, 47*). A possible explanation for the limited effects of A*β* therapies on AD progression is that intervention started too late in the AD timeline. This is consistent with the dATN model, which predicts that A*β* reduction before there is tau acceleration is necessary for optimal intervention with A*β* targeting therapies, since after this time, tau can accumulate faster and cause neurodegeneration.

Stopping A*β* after tau acceleration will result in slowing of disease progression (as observed in clinical trails), whereas reducing A*β* before tau acceleration will halt the progression of AD into secondary tauopathy. Here, we have introduced a spatial pharmacokinetic model and illustrate how whole-brain modelling may inform the effectiveness of different administration methods and protocols. Existing PKPD modelling of AD relies on compartmental models of A*β*, tau aggregation and summary levels of pathology, such as the centiloid scale (*45, 48, 49*). We provide a novel method of modelling the effectiveness of drugs in vivo that may be useful for clinicians and pharmaceutical scientists in designing and conducting clinical trials. Future work should aim to incorporate such models with clinical trial data to enhance the fidelity with which drug effects are observed and quantified.

Additionally, this work may have implications for future tau targeting therapies. We and others have previously demonstrated that tau spread is dominated by production, not transport, in *A*^+^*T*^+^ individuals (*27, 69*). However, in this work, we show that the dynamics tau in the *A*^+^*T*^+^ group are neither production-dominated or transport-dominated, since both parameters are of the same magnitude. We can see from the integrative approach taken here that the production dominated effect observed by Meisl et al. 2021 (*69*) and Chaggar et al. 2025 (*27*) stems from the accelerating impact of A*β* on tau, which in turn accelerates neurodegeneration. This suggests an effective strategy for intervention would be to limit the interactions of A*β* and tau, to prevent tau for entering a production-dominated phase. This may be partly achieved with A*β* reducing therapies, but also with putative tau targeting therapies. For example, tau therapies that aim to limit the extracellular spread of tau (*70*) could be used early in AD to prevent widespread deposition of tau seeds to cortical areas of high A*β* load. Alternatively, tau targeting therapies aimed at reducing intracellular tau (*71*) could be used for patients who have more advanced AD. In this case, our model suggests that intracellular tau reduction therapies should be used in conjunction with A*β* reducing therapies to maximise their effect, otherwise anti-tau therapies will be counteracted by the accelerating effect of A*β*.

Although the work here advances on previous modelling work (*27, 44, 72*), we still face similar limitations. First, we still rely on fixing important parameters, such as baseline and carrying capacities of A*β* and carrying capacities in PART. Fixed parameters for A*β* are likely to be robustly identified from cross-sectional analysis given the abundance of A*β* PET data, however, by fixing their values we ignore potential inter-subject variability and any dynamical changes over time. In estimating the PART carrying capacities, we are limited by the number subjects in whom PART can be detected in the absence of A*β*, namely the *A*^−^*T*^+^ ADNI cohort. Additionally, by fixing these values we are unable to explain how heterogeneities in A*β* deposition and early tau burden occur, and if or how they change over time. The importance of A*β* heterogeneity is marked by recent studies demonstrating the existence of A*β* spatiotemporal subtypes (*73*) that overlap with spatial subtypes of tau and cortical neurodegeneration (*74*–*76*). Future work should examine whether A*β* deposition is predictive of tau deposition across spatial subtypes and whether correlations between parameters exist that can be used to further characterise subtypes. Such information could be included to provide more constrained prior information on parameters based on individual A*β* deposition to allow for more personalised predictions for limited longitudinal data. Furthermore, the model reduction we perform comes at the cost of losing mechanistic insight. To address more specific questions about interaction mechanism, we require more detailed modelling of A*β*/tau interactions, cellular transport and interactions between oligomers of different sizes. Such modelling could be applied to animal models and future work might benefit from addressing not only macroscale modelling applied to multimodal human neuroimaging data but also microscale modelling of animal and in-vivo models. Another major limitation of the current study is the limited sample size that limits conclusions about population-level dynamics. Despite this, we note that posterior densities for individual parameters are generally identifiable, showing little variation. Therefore, while individual parameters are unlikely to change dramatically provided more data, population-level parameters may change. In the absence of more data, future work should validate the results presented here on other longitudinal datasets. In addition to these limitations, we note the heavy data burden required to calibrate and apply the ATN model that will greatly limit its utility in clinical practice, where longitudinal PET data is rarely available. However, such data is routinely collected for clinical trials and pharmaceutical research and the dATN model may provide useful applications for patient selection and pharmacodynamic modelling. We note also that for the colocalisation and PKPD analysis, we use parameters derived from an *A*^+^*T*^+^ group, which may not translate to the early tau group if there are time-dependent changes in parameters or changes to initial conditions. This is partly addressed in Fig. S3, where we reproduce Fig. 4F using varying centiloid thresholds and note some sensitivity to the initial conditions. Namely, in the group with the lowest centiloid threshold, with a mean CL of 35, the colocalisation point occurs after the inflection point, while for higher centiloid thresholds, there is close correspondence between colocalisation and the inflection point. This effect is likely due to the poor identification of cortical tau seeds with PET in early AD (*68*). This limitation should be considered when applying the dATN model to early AD groups.

Our analysis provides further evidence for a mechanistic link between A*β* and tau. In sum, we show that A*β* catalyses the local production of tau, pushing tau into an accelerated production-dominated regime, in which tau seeds amplify in a regionally heterogenous manner related to local A*β* deposition. This relationship between A*β* and tau is clinically meaningful, and we show how colocalisation between A*β* and tau may be an important clinical biomarker for predicting interventional success through A*β* targeting therapies. More broadly, our results rely on a general modelling and simulation framework that can be utilised and extended to investigate AD pathology in a mechanism-based manner.

## MATERIALS AND METHODS

### Study Design

The goal of this study was to investigate interactions between Aβ, tau, neurodegeneration by pairing a dynamical model of ATN biomarker with neuroimaging data. More specifically, we aimed to identify whether regional Aβ load drives regional tau heterogeneity, if regional tau load drives neurodegeneration, and how Aβ targeting therapies may affect downstream regional biomarker progression. To do so, we use the maximum amount of data available from ADNI and BF2 that met a predetermined selection criterion: that individuals had at least 2 Aβ PET scans and at least 3 tau PET scans. No subjects were excluded as outliers from the analysis. Inference from the model and data was performed multiple times using different starting conditions to ensure reliable convergence. Data were not randomised, and the investigators were not blinded to the individuals’ diagnoses or biomarker profiles. The analysis of the effects of Aβ targeting therapies on regional biomarker progression was conducted in silico; therefore, no treatments were administered to real individuals.

### Demographics

Throughout this study we use neuroimaging data available through the Alzheimer’s Diseaes Neuroimaging Initiative (ADNI) (adni.loni.usc.edu) and the BioFINDER-2 study (NCT03174938) (71). ADNI is a public-private partnership with the aim of using serial biomarkers to measure the progression of AD. For up-to-date information, see www.adni-info.org.

We make use of longitudinal Aβ PET, tau PET and structural MRI from both cohorts. We use data for two purposes: (1) first to determine the fixed model parameters; (2) post-hoc analysis (colocalisation and PKPD analysis) with data-derived initial conditions for Aβ and tau PET in an early tau group. To determine regional Aβ values, we use Aβ PET data from all available subjects with at least two scans ADNI. To determine carrying capacities for PART pathology, we use all available tau PET scans for *A*^−^ subjects only. For inference, we use *A*^+^*T*^+^ subjects with at least two Aβ PET scans and’ three tau PET scans. For colocalisation and PKPD modelling, we use an early tau group defined by a mean CL ≥ 40, when early tau presence is detectable with PET (*51, 52*). This is achieved with a threshold of 70 CL in ADNI and 72 CL in BF2.

**Table 4.1.** Demographics for ADNI and BF2 cohorts.

|  | ADNI |  | BF |  |
| --- | --- | --- | --- | --- |
| Variable | Inference | Early tau | Inference | Early tau |
| N. Subjects | 34 | 69 | 48 | 325 |
| Age (years) | 71.65 | 71.71 | 70.587 | 73.00 |
| Gender (% Female) | 0.52 | 0.52 | 0.438 | 0.51 |
| Education (years) | 16.7 | 16.8 | 12.052 | 12.582 |
| CN (%) | 0.26 | 0.53 | 0.14 | 0.25 |
| MCI (%) | 0.50 | 0.40 | 0.86 | 0.70 |
| AD (%) | 0.24 | 0.07 | 0.000 | 0.05 |
| CL | 84 | 41 | 73 | 40 |
| CL std | 34 | 19 | 32 | 19 |

### Data Processing

All PET and MRI data from ADNI was downloaded as preprocessed derivatives summarised on the FreeSurfer Deskian-Killiany (DK) atlas. For FBB and FBP Aβ PET, we calculate the standardised uptake volume ratio (SUVR) using the composite reference region provided by ADNI. A subject is deemed to be A^+^ if their composite FBB SUVR > 0.78 or FBP SUVR > 0.74 in the ADNI cortical summary region for all scans. FTP tau PET was normalised using the inferior cerebellar reference region. Tau positivity is determined using the average SUVR in two composite regions, a medial temporal composite comprising the bilateral entorhinal cortex and amygdala, and a neocortical composite comprising the bilateral inferior and middle temporal lobes, as detailed in (*27, 35*). A subject is said to be T^+^ if either their medial temporal SUVR > 1.375 or cortical SUVR > 1.395 in their most recent PET scan and T^−^ otherwise.

Data from BF2 uses FMM for Aβ PET and RO948 for tau PET. All participants were recruited at Skåne University Hospital and the Hospital of Ängelholm, Sweden. All details about the cohort have been described previously (*77*). Amyloid positivity required FMM SUVR > 1.03 with using a cerebellar reference region (*35*). The tau PET analysis pipeline is detailed in (*77*). SUVR images were generated using the inferior cerebellum as a reference region, and average SUVR was extracted for regions in the DK atlas. Tau positivity is determined in the same way as for ADNI data, with medial temporal SUVR cutoff > 1.248 and cortical SUVR > 1.451.

We use dMRI data from the Human Connectome Project (*78, 79*) to derive a normative connectome from young healthy individuals that we use to model the transport of tau along axonal connections. The diffusion weighted MRI images of 150 individuals in the Human Connectome Project were processed using the probtrackx program in FSL (*80*), with 10000 random streamline samples from a sphere surrounding the center of each voxel. The streamlines are summarised on the DK atlas, comprising 68 cortical regions, in addition to the bilateral Amygdala and hippocampus. Therefore, there are R = 72 regions in total. We have an adjacency matrix A ∈ ℝ^R×R^ which defines the connectome on the DK atlas. which we normalise by its maximum value, 𝒜_*ij*_ = A_*ij*_/max_*ij*_A, so values lie in the [0,1] interval. Finally, entries below a threshold of 0.01 are excluded from the matrix to reduce the occurrence of spurious connections inferred during tractography. The graph Laplacian, L is used to model the transport of tau along axonal connections and is defined as **L** = diag(**𝒜 ⋅ 1**) − **𝒜**.

### Mechanistic Modelling

We start with a modified version of the network heterodimer model that incorporates A*β*-tau interactions (*43*). For variables *u*_*i*_ = *u*_*i*_(*t*) and ũ_*i*_ = ũ_*i*_(*t*) representing toxic and healthy A*β* concentration, respectively, and *p*_*i*_ = *p*_*i*_(*t*) and 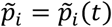 healthy and toxic tau concentration, respectively, at each node *i* = 1, …, *R*, the coupled model is:

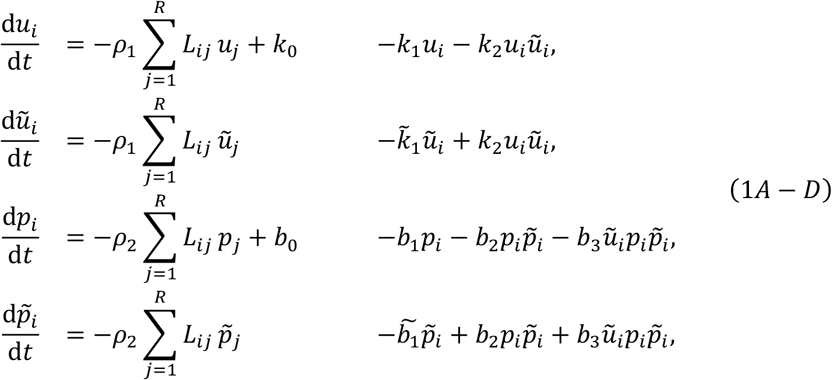

with *u*_*i*_(0) = *u*_*i*,0_, ũ_*i*_(0) = ũ_*i*,0_, *p*_*i*_(0) = *p*_*i*,0_, and 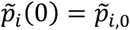 for *i* = 1, …, *R*. For A*β*, there are rate parameters for the production and clearance of healthy A*β, k*_0_ and *k*_1_ respectively, clearance of toxic A*β*, 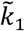, and conversion of healthy to toxic A*β* at rate *k*_2_. Similarly, for tau, there are rate parameters for the production and clearance of healthy tau, *b*_0_ and *b*_1_, respectively, clearance of toxic tau, 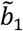, conversion of healthy to toxic tau, *b*_2_, and conversion of healthy to toxic tau catalysed by A*β, b*_3_. For healthy and toxic A*β* and tau, transport along the connectome is described by the graph Laplacian, **L**, with transport rates *ρ*_1_ and *ρ*_2_, for A*β* and tau, respectively. While the model is informative about the mechanisms of AD proteopathy, there is insufficient information in neuroimaging data from which to calibrate the numerous parameters. Therefore, we simplify it to obtain a model that captures essential features of the system but has with fewer parameters that can be calibrated with existing A*β* and tau PET data. We follow a similar procedure to previous work (*81*).

We assume that the transport of A*β* is negligible across the brain network, following evidence from cross-sectional analysis showing that A*β* is likely to be densely deposited around the cortex during early stages of AD (*18*) and the homeostatic level of healthy A*β* do not affect the toxic levels of A*β*. Therefore, Equations (1A–1B) simplify to a single equation to describe the evolution of toxic A*β*, as in (*27*):

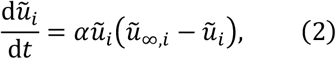

for *i* = 1, …, *R*, where 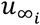 is the regional carrying capacity for A*β*.

To linearise (1C–1D), we assume a healthy and homogenous state, then rewrite Equation 1C to find *p* as a function of 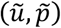,

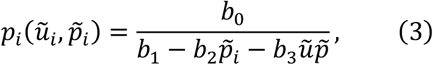

that we expand to first order to obtain,

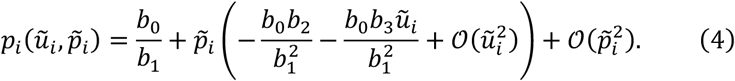

Substituting this last expression into Equation 1D, we obtain the following expression

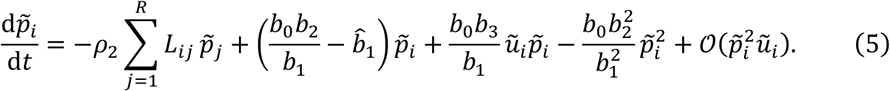

Following the assumption of a healthy state, we drop higher-order terms of toxic species, since they will be small relative to quadratic terms. Then, grouping like powers of 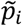, we obtain

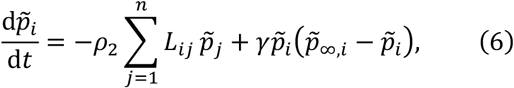

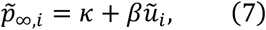

where

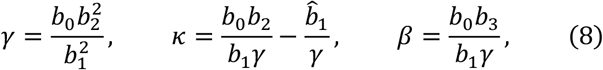

and 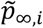 is the regional carrying capacity in the absence of transport. This nonlinear model is based on a linear approximation of certain species appearing in Equation 1.

To develop a full model of the ATN pathway, we introduce a simple model of neurodegeneration, with variable N_*i*_ = N_*i*_(*t*) for *i* = 1, …, *R*, following the assumption that neurodegeneration is primarily correlated with tau (*38, 39*). Therefore, the full ATN model is given by a system of 3*R* equations in the 3*R* variables 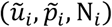 for *i* = 1, …, *R*,

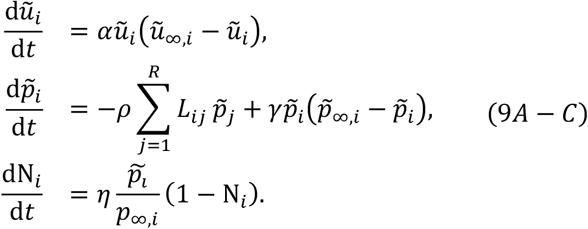

Where 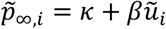 is the carrying capacity for tau. We note that it has a non-A*β* dependent part and an A*β* dependent part. Note that we drop subscript notation for *ρ*, since we neglect the transport effect on A*β*.

To integrate longitudinal neuroimaging data, we must specify a measurement model that connects the amount of toxic proteins *u*_*i*_ and *p*_*i*_ for *i* = 1, …, *R* to the SUVR values. Hence, we must account for regional baseline values of A*β* and tau PET SUVR that would be present in the absence of the disease. Therefore, we extend Equations 9A-C to a model that shifts the baseline values of A*β* and tau PET through the variables

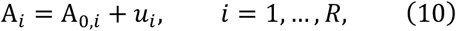

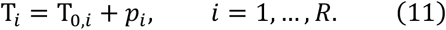

where, A_*i*_ and T_*i*_ represent the A*β* and tau SUVR at the *i*-th region, and A_0,*i*_ and T_0,*i*_ are baseline SUVR values at the *i*-th region. Incorporating these changes into Equation 9, we have for *i* = 1, …, *R*:

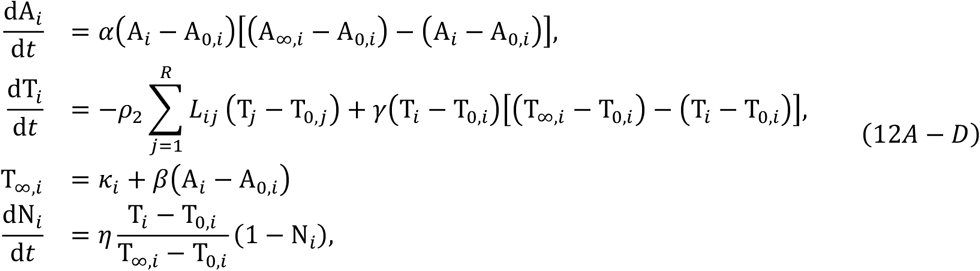

with A_*i*_(0) = A_*i*,0_, T_*i*_(0) = T_*i*,0_, N_*i*_(0) = N_*i*,0_, and where, T_0,*i*_ is the regional baseline value for tau, *κ*_*i*_ is the regional tau carrying capacity in the absence of A*β*, T_∞,*i*_ is the regional tau carrying capacity in the presence of A*β*, A_0,*i*_ and A_∞,*i*_ are the baseline values and carrying capacities for A*β* SUVR. Note that we further assume **κ**, representing the natural vulnerability for tau accumulation, of varies regionally, consistent with non-pathological tau accumulation seen in PART.

For colocalisation and PKPD modelling we use a scaled version of this model to the unit range. By applying the change of variables:

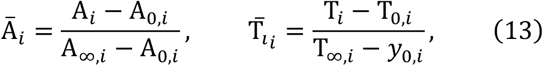

for *i* = 1, …, *R*, we retrieve the following system of equations for A*β* concentration, tau concentration and neurodegeneration, 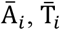 and N_*i*_, respectively, evolving in [0,1].

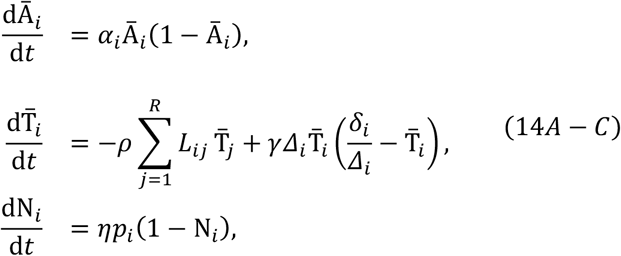

where *α*_*i*_ = *α*(A_∞,*i*_ − A_0,*i*_), 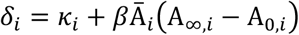 and *Δ* = *κ*_*i*_ + *β*(A_∞,*i*_ − A_0,*i*_). Note that as *t* → ∞ and for 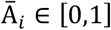 and *α* > 0, then *δ*_*i*_ → *Δ*_*i*_. To ensure 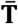 remains in the unit interval, we neglect the (small) asymmetry in **L** induced by this change of variables.

### Estimating Fixed Model Parameters

There are four parameters we estimate from cross-sectional data, baseline A*β* and tau SUVR values **x**_0_, **y**_0_, carrying capacities for A*β* PET, **x**_∞_, and the PART SUVR carrying capacities **κ**. To estimate **x**_0_ and **x**_∞_ we reproduce the analysis of Jack et al. (*82*) and Whittington et al. (*18*) to temporally order subjects with at least two scans according to their A*β* load. For ADNI and BF2, we use the Berekely summary composite region SUVR (*83*) in *N* = 205 ADNI subjects with FBB scans, *N* = 763 ADNI subjects with FBP scans, and *N* = 813 BF2 subjects with FMM scans. We estimate parameter vectors for each tracer separately. Using the SUVR summary data, we approximate the derivative at each point by calculating the forward difference between an individual’s baseline and final scan.

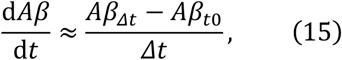

where *Aβ*_*t*0_ is the initial A*β* load for a subject, *Δt* is the time between initial and final scan, and *Aβ*_*Δt*_ is the A*β* load at the final scan. The 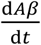 data are binned to intervals of 0.05 SUVR to smooth the noisy derivative estimates and second-degree polynomial is fit by minimising the least squares error. The polynomial is integrated to retrieve a logistic curve and individuals are assigned a *time* on this curve corresponding to their baseline A*β* level. A sigmoid curve is fit to the SUVR values for each region in the DK atlas, using the subject-level SUVR and A*β*-time estimates to order the SUVR. The sigmoid function is given as

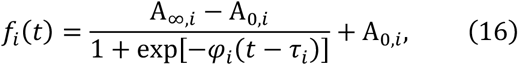

where *φ*_*i*_ is a regional production rate, *τ*_*i*_ is the time at which the half-maximal concentration is reached, A_0,*i*_ is regional baseline value, and A_∞,*i*_ is the regional carrying capacity. This model was fit to each region by minimising the least squares error.

To determine the PART SUVR carrying capacities, **κ**, we apply a Gaussian mixture modelling approach (*26, 27*) to 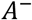 subjects from ADNI (*N* = 582) and BF2 (*N* = 943), determined using the SUVR thresholds detailed above. We restrict our analysis to temporal regions only, since these are regions affected by PART (*28*). We fit a one-component and a two-component Gaussian mixture model to cross-sectional regional SUVR data and their model fit is compared using the Akaike information criteria (AIC) (*26*). Regions are considered *T*^+^ if the AIC for the two-component model is lower than the one-component model. For regions that are *T*^+^, the PART carrying capacity is then given by the 99th percentile of the *T*^+^ distribution. If a one-component Gaussian mixture model for a region provides a superior fit, the regional PART carrying capacity is equal to the regional baseline SUVR value.

### Inference with ATN Biomarkers

We use longitudinal A*β* PET, tau PET and sMRI from *A*^+^*T*^+^ subjects with at least two A*β* scans and at least three tau scans to calibrate the dATN model. We take average SUVR values for regions defined by the DK atlas, with the addition of the bilateral amygdala and hippocampus, resulting in *R* = 72 regions. To ensure numerical stability when solving the dATN model, regional A*β* PET data denoted **Y**_A_ are normalised to lie between the baseline and carrying capacities. Regional tau PET data, denoted **Y**_T_, is normalised to be above the regional baseline tau values. Regional volumetric data, 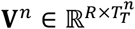 for *n* = 1, …, *N*, are provided with each PET scan and these are used to measure regional neurodegeneration, **Y**_N_. Regional volumes are normalised to their baseline total intracranial volume. Then, all volume measures for an individual divided by their values at the baseline scan and neurodegeneration is measured as 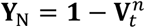 for 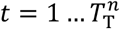 and *n* =1, …, *N*, where 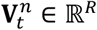 is the *t*th scan for the *n*th subject. Therefore, at the initial scan for each subject, the measured neurodegeneration per region is 0 and increases as brain volumes decrease.

In total, we have *N* = 34 *A*^+^*T*^+^ subjects in ADNI and *N* = 48 in BF2. For a given cohort, the full dataset is denoted 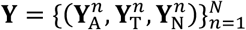, where subscripts A, T and N are used to denote biomarker data for each of the ATN biomarkers. Each subject has 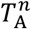 A*β* PET scans, and 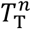 tau PET scans and sMRI scans, for *n* = 1, …, *N*. Note that we use the corresponding sMRI scans obtained with tau PET scans. Subjects may have different measurement times for A*β* PET scans and tau PET scans. Therefore, we have 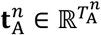 and 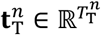 for *n* = 1, …, *N* subjects A*β* PET and tau PET scan times, respectively. Additionally, we denote the initial conditions as the vector 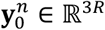, comprising the initial A*β* PET, tau PET and sMRI values for the *n*th subject. For the *n*th subject the data generating function is:

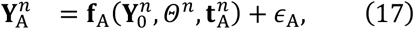

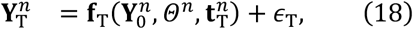

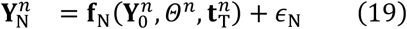

where **f**_A_, **f**_T_, **f**_N_ are solutions of ATN model, Equations 12A-D, respectively, 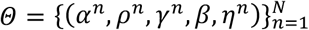 are the model parameters for the *n*th subject, and *ϵ*_A_, *ϵ*_T_, *ϵ*_N_ represent measurement noise process specific to each biomarker. We pool *β* across individuals to aid identifiability. We assume that each biomarker has independent and identically distributed Gaussian noise, parametrised by **σ** = {*σ*_A_, *σ*_T_, *σ*_N_}. Our generative model for the *n*th subject and each biomarker is then:

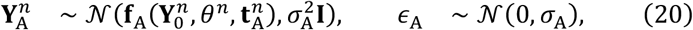

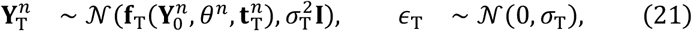

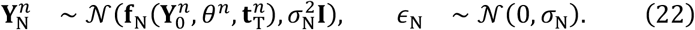

We extend this model to a hierarchical Bayesian model to explicitly model inter-individual differences by introducing population-level parameters *Ω* = {α_*μ*_, *α*_*σ*_, *ρ*_*μ*_, *ρ*_*σ*_, *γ*_*μ*_, *γ*_*σ*_, *η*_μ_, *η*_σ_}, upon which each of the respective model parameters depend. The log likelihood for a single subject under the hierarchical model is given by:

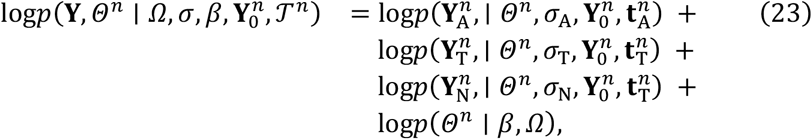

where 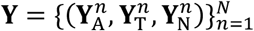 is the collection of all ATN biomarker data in the cohort. The first three terms represent the log-likelihood contribution for each of the ATN components in the model given their respective longitudinal data, and the last term represents the contribution from the population-level model. This can then be extended to the full cohort of *A*^+^*T*^+^ subjects by accumulating the log likelihood. The full posterior is then given by:

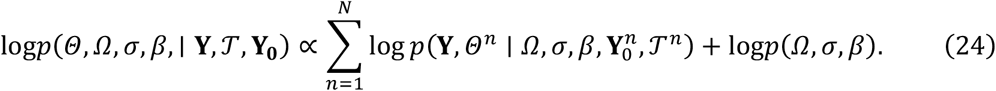

We place weakly informative priors on the model parameters, provided in Table 2. For model parameters that are strictly positive, we use standard Lognormal priors as continuous positive distributions skewed toward smaller parameter values that produce dynamics on the scale of years. For the coupling parameter *β*, we use a half-Normal prior of 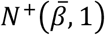, where 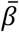 is the coefficient of a linear model fit to A*β* and tau carrying capacities, for each combination of A*β* and tau tracer used (FBB/FTP, FBP/FTP, FMM/RO948).

**Table 2.** Prior distributions for hierarchical model parameters.

| Parameter | Description | Prior | Support |
| --- | --- | --- | --- |
| Population-level parameters |  |  |  |
| $\alpha_\mu$ | Aβ production mean | Lognormal(0,1) | $[0, \infty]$ |
| $\alpha_\sigma$ | Aβ production SD | $\mathcal{N}^+(0,1)$ | $[0, \infty]$ |
| $\rho_\mu$ | tau transport mean | Lognormal(0,1) | $[0, \infty]$ |
| $\rho_\sigma$ | tau transport SD | $\mathcal{N}^+(0,1)$ | $[0, \infty]$ |
| $\gamma_\mu$ | tau production mean | Lognormal(0,1) | $[0, \infty]$ |
| $\gamma_\sigma$ | tau production SD | $\mathcal{N}^+(0,1)$ | $[0, \infty]$ |
| $\eta_\mu$ | neurodegeneration rate mean | Lognormal(0,1) | $[0, \infty]$ |
| $\eta_\sigma$ | neurodegeneration rate SD | $\mathcal{N}^+(0,1)$ | $[0, \infty]$ |
| Subject-level parameters |  |  |  |
| $\alpha_i$ | Aβ production | $\mathcal{N}^+(\alpha_\mu, \alpha_\sigma)$ | $[0, \infty]$ |
| $\rho_i$ | tau transport | $\mathcal{N}^+(\rho_\mu, \rho_\sigma)$ | $[0, \infty]$ |
| $\gamma_i$ | tau production | $\mathcal{N}^+(\gamma_\mu, \gamma_\sigma)$ | $[0, \infty]$ |
| $\beta_{fbb}$ | Aβ/tau coupling (FBB/FTP) | $\mathcal{N}^+(3.2, 1.0)$ | $[0, \infty]$ |
| $\beta_{fbp}$ | Aβ/tau coupling (FBP/FTP) | $\mathcal{N}^+(3.6, 1.0)$ | $[0, \infty]$ |
| $\beta_{fmm}$ | Aβ/tau coupling (FMM/RO948) | $\mathcal{N}^+(2.3, 1.0)$ | $[0, \infty]$ |
| $\eta_i$ | neurodegeneration rate | $\mathcal{N}^+(\eta_\mu, \eta_\sigma)$ | $[0, \infty]$ |
| Noise parameters |  |  |  |
| $\sigma_A$ | Aβ PET observation error | $\Gamma^-(2,3)$ | $[0, \infty]$ |
| $\sigma_T$ | tau PET observation error | $\Gamma^-(2,3)$ | $[0, \infty]$ |
| $\sigma_N$ | sMRI observation error | $\Gamma^-(2,3)$ | $[0, \infty]$ |

To sample from the posterior distributions we use a NUTS algorithm with an acceptance rate of 0.8 and a dense Euclidean metric (*84, 85*). We collect four chains starting from different initial parameters with 1000 samples each. For all chains we observe no numerical errors associated with the Hamiltonian integration and 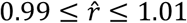, showing the chains converged.

### A*β*/tau Colocalisation

The dATN model (Equation 14) and posterior samples are applied to subjects in ADNI and BF2. For initial conditions, we use the mean FBB and FTP SUVR for regions in the DK atlas from *N* = 69 subjects with from an early tau group, defined as having a *CL* ≥ 40, where there is certain A*β* pathology and the emergence of tau pathology (*51, 52*). To threshold the data, we use the SUVR value *μ*_i_ + *σ*_i_ for, where *μ*_i_ and *σ*_i_ are the mean and standard deviation of healthy tau SUVR signal for each region *i*, estimated using a two-component Gaussian mixture model applied to cross-sectional FTP and RO948 data separately. Signal below this threshold is set to zero and the SUVR values are scaled to concentrations using the transformations in Equation 13. For both A*β* and tau, we symmetrise the initial conditions across hemispheres by setting the regional signal to the average of the two regions. The thresholds for colocalisation are determined using the velocity and jerk of A*β* and tau respectively. For the solution to the ODE Equation 14, *f*^A^ and *f*^T^ for A*β* and tau concentration, respectively, with initial conditions given by the mean A*β* and tau from early tau cohort and the mean individual posterior parameters, we define the regional threshold A*β* concentration saturation as

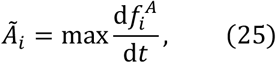

and the regional threshold for tau concentration as

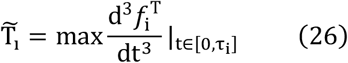

where *τ*_*i*_ is the time at which 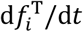 is maximised for *i* = 1 … *R*. Then, a region is considered to have undergone colocalisation if the A*β* and tau concentrations are both greater than their regionally defined thresholds. To calculate the colocalisation order and probability, we simulate progression using the initial conditions derived from averages A*β* PET and tau PET scans from the early tau cohort and posterior samples from the pooled model. To calculate the *colocalisation order*, we use the mean individual-level posterior parameters. To calculate the *initial colocalisation probability* we iterate over all samples from the posterior distribution and identify the percentage of samples that lead to a given region being the site of initial concentration.

### Intervention Modelling

For the PK model, we only model the drug concentration in the central compartment (the brain), entering through regions around the major blood arteries surrounding the lateral ventricles. We denote this set of regions as *V* and they comprise the following regions of the DK atlas: rostral anterior cingulate, caudal anterior cingulate, posterior cingulate, isthmus cingulate, hippocampus, and amygdala. We model drug administration using an exponential decay model for the *i*-th region as follows,

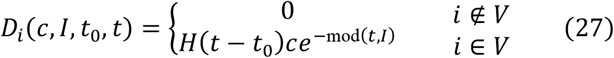

where *D*_i_ describes the regional drug administration at time *t, c* ∈ ℝ^+^ is the drug concentration, *I* ∈ ℝ^+^ is the dosing interval in months, and *t*_0_ ∈ ℝ^+^ is the initial administration time and *H* is the Heaviside function.

We model drug entry to regions not in *V* through by diffusion of drug using the graph Laplacian of a distance matrix for the DK atlas, denoted by 𝒟, and drug diffusion coefficient *ρ*_m_. 𝒟 is constructed from an adjacency matrix 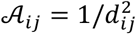, where *d*_*ij*_ is the distance between regions *i* and *j*. We also model drug elimination from the brain with rate constant *λ*, and a linear damping by drugs on A*β* with rate *ζ*. The carrying capacity of tau, which depends on regional A*β* concentration, is assumed to remain constant upon A*β* clearance. The full model of drug concentration at the *i*-th region coupled with the ATN model is then:

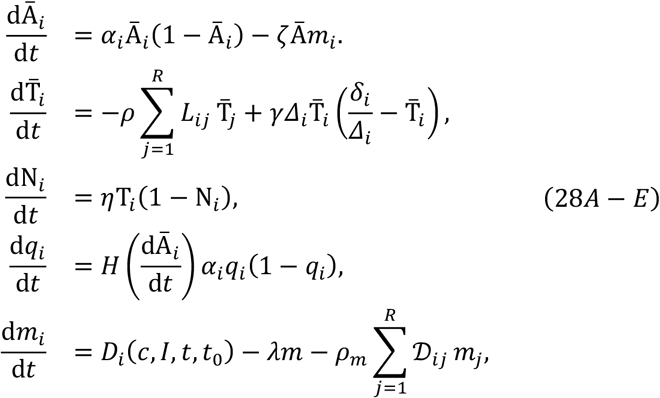

Where *H* is the Heaviside function, *q*_*i*_ is a new variable that follows the trajectory of A*β* while 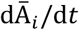 is positive, and is 0 otherwise, and *δ*_*i*_ = *κ*_*i*_ + *β*q_*i*_(A_∞,*i*_ − A_0,*i*_), modelling the assumption that the regional tau carrying capacity remains constant after A*β* clearance. The dATN-PKPD model is then solved in the left hemisphere using average initial conditions from an early tau cohort in ADNI, who have at least one FBB and FTP scan (see Table 1 for demographic details), and the average individual-level ATN model parameters rescaled to months and PKPD parameters in Table 3. We simulate the ATN-PKPD model over a 30-year period, with different drug administration starting points of *t*_0_ ∈ {2*x* ∣ 0 ≤ *x* ≤ 15, *x* ∈ ℕ} and continuing administration every month. Initial conditions for A*β* and tau are derived from FBB and FTP PET, respectively, in the early tau group, using mean SUVR values for regions in the DK atlas. Initial conditions for neurodegeneration, q_’_, and drug concentration, *m*_i_ are 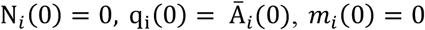, respectively for *i* = 1, …, *R*. From these simulations, we calculate endpoint tau and neurodegeneration as their final simulated values in Braak stage 1-3 regions at *t* = 30. Then relative change in endpoint tau and neurodegeneration is calculated as the difference between endpoint values for simulations starting at *t*_0_ = *n* and *t*_0_ = *n* − 2 for n ∈ {2*x* ∣ 1 ≤ *x* ≤ 15, *x* ∈ ℕ}.

**Table 3.** Model parameters used to simulate from the dATN-PKPD model.

| Parameter | Description | Value |
| --- | --- | --- |
| $\alpha$ | Aβ production | 0.031 / month |
| $\rho$ | tau transport | 0.006 / month |
| $\gamma$ | tau production | 0.011 / month |
| $\beta$ | Aβ/tau coupling | 4.864 |
| $\eta$ | Neurodegeneration rate | 0.013 / month |
| $\rho_m$ | drug diffusion | 0.125 / month |
| $\zeta$ | drug effect | 0.01 / month |
| $c$ | dose concentration | 400.000 mg |
| $\lambda$ | drug clearance | 0.500 |
| $I$ | dosing interval | 1 / month |

### Statistics

We use a Bayesian hierarchical model and the NUTS algorithm to perform inference from the dATN model and biomarker data. For ADNI and BF2, we sampled 4 chains starting from different initial conditions, each with 1000 samples. Chain convergence was assessed using the 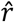 value with the condition that 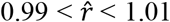. Population-level distributions are shown as histograms, while individual-level distributions (Fig. S2) are shown as kernel density estimates to improve visibility. The fit between the dATN model predictions and observations is assessed using the root-mean-squared error and *R*^2^ values with a 95% confidence interval.

## Acknowledgments

Data used in preparation of this article were obtained from the Alzheimer’s Disease Neuroimaging Initiative (ADNI) database (adni.loni.usc.edu). As such, the investigators within the ADNI contributed to the design and implementation of ADNI and/or provided data but did not participate in analysis or writing of this report. A complete listing of ADNI investigators can be found at: http://adni.loni.usc.edu/wp-content/uploads/how_to_apply/ADNI_Acknowledgement_List.pdf.

Data collection and sharing for this project was funded by the Alzheimer’s Disease Neuroimaging Initiative (ADNI) (National Institutes of Health Grant U01 AG024904) and DOD ADNI (Department of Defense award number W81XWH-12-2-0012). ADNI is funded by the National Institute on Aging, the National Institute of Biomedical Imaging and Bioengineering, and through generous contributions from the following: AbbVie, Alzheimer’s Association; Alzheimer’s Drug Discovery Foundation; Araclon Biotech; BioClinica, Inc.; Biogen; Bristol-Myers Squibb Company; CereSpir, Inc.; Cogstate; Eisai Inc.; Elan Pharmaceuticals, Inc.; Eli Lilly and Company; EuroImmun; F. Hoffmann-La Roche Ltd and its affiliated company Genentech, Inc.; Fujirebio; GE Healthcare; IXICO Ltd.; Janssen Alzheimer Immunotherapy Research and Development, LLC.; Johnson and Johnson Pharmaceutical Research and Development LLC.; Lumosity; Lundbeck; Merck and Co., Inc.; Meso Scale Diagnostics, LLC.; NeuroRx Research; Neurotrack Technologies; Novartis Pharmaceuticals Corporation; Pfizer Inc.; Piramal Imaging; Servier; Takeda Pharmaceutical Company; and Transition Therapeutics. The Canadian Institutes of Health Research is providing funds to support ADNI clinical sites in Canada. Private sector contributions are facilitated by the Foundation for the National Institutes of Health (www.fnih.org). The grantee organization is the Northern California Institute for Research and Education, and the study is coordinated by the Alzheimer’s Therapeutic Research Institute at the University of Southern California. ADNI data are disseminated by the Laboratory for Neuro Imaging at the University of Southern California.

Data were provided (in part) by the Human Connectome Project, WU-Minn Consortium (Principal Investigators: David Van Essen and Kamil Ugurbil; 1U54MH091657) funded by the 16 NIH Institutes and Centers that support the NIH Blueprint for Neuroscience Research; and by the McDonnell Center for Systems Neuroscience at Washington University.

The precursor of 18F-flutemetamol was sponsored by GE Healthcare.The precursor of 18F-RO948 was provided by Roche.

## Funding

Engineering and Physical Sciences Research Council EP/L016044/1 (PC)

Engineering and Physical Sciences Research Council EP/R020205/1 (AG)

SciLifeLab and Wallenberg Data Driven Life Science Program KAW 2020.0239 (JWV)

and European Research Council StG-101221737 (JWC)

Wellcome Trust Collaborative Award 215573/Z/19/Z (SJ)

Wellcome Senior Research Fellowship 221933/Z/20/Z (SJ)

National Science Foundation grant NSF 2325276 (TT)

National Institute of Aging R01AG083740

Alzheimer’s Association 22AARF-1029663, ZEN24-1069572

The Alzheimer’s Association in collaboration with the GHR Foundation ALZSI-26-1523522

The European Research Council ADG-101096455

European Commission/Horizon Europe 101108819, 101153323

ERA PerMed ERAPERMED2021-184

The Michael J Fox Foundation MJFF-025507, MJFF-025741

Global Research Platform LLC

The Cure Alzheimer’s Fund F2025/1066

Swedish Research Council 2021-02219, 2022-00775, 2023-06428

The Swedish Alzheimer Foundation AF-981132, AF-980907, AF-968598, AF-1011949, AF1011799

The Swedish Brain Foundation FO2023-0163, FO2024-0133-HK-46, FO2024-0284, FO2024-0385, FO2025-0055

The Swedish Parkinson Foundation 1485/23, 1589/24

The Strategic Research Area MultiPark (Multidisciplinary Research in Parkinson’s disease) at Lund University WASP and DDLS Joint call for research projects WASP/DDLS22-066

The Skåne University Hospital Foundation Regional research support 2025-2024-2925, 2025-2024-2862

The Swedish federal government under the ALF agreement 2022-Project0080, 2022-Project0107, 2022-Project0062, 2022-YF0017, 2024-YF0048, 2024-ST0026

Lilly Research Award Program F2025/932

Avid Pharmaceuticals

F Hoffman-La Roche AG F2025/453

Biogen

Bristol Myers Squibb

CellInvent F2025/1321

Eisai

Fujirebio

GE Healthcare

The Knut and Alice Wallenberg Foundation 2017-0383

Konung Gustaf V:s och Drottning Victorias Frimurarestiftelse

The Kamprad Foundation 20243058

Bundy Academy

Greta and Johan Kock Foundation F2024/228, F2024/2330, F2024/197, F2025/160, F2024/198, F2024/2332, F2025/319

The Crafoord Foundation 20240888

The Rönström Family Foundation AF1011799, FRS 0013, FRS-0003, FRS-0004, AF-1011949

The Royal Physiographic Society F2025/1092

Stiftelsen för Gamla Tjänarinnor 2024-250

Thorsten and Elsa Segerfalk’s Foundation F2024/1053, F2025/1320

Fredrik and Ingrid Thuring’s Foundation 2024-099

## Author contributions

Conceptualization: PC, JWV, RA, GK, NMC, OH, AG

Methodology: PC, JWV, RA, GK, NMC, OH, AG

Formal Analysis: PC, AG

Investigation: PC, JWV, NMC, OH, AG

Data Curation: OS, ES

Visualization: PC, JWV, NMC, OH, AG

Funding acquisition: PC, JWV, RO, SP, NMC, OH, AG

Supervision: SM, SJ, GK, NMC, OH, AG

Writing – original draft: PC, JWV, NMC, OH, AG

Writing – review & editing: SBB, DLA, PRB, JLS, EH

## Competing interests

P.C. has received consultancy fees from Roche. NMC has received speaker/consultancy fees from Biogen, BioArctic, Eli Lilly, Merck, Novo Nordisk, and Owkin. OH is an employee of Lund University and Eli Lilly.

## Data and materials availability

Pseudonymized data from ADNI are available upon application through adni.loni.usc.edu. Pseudonymized data from the BioFINDER study can be shared by request from a qualified academic as long as data transfer is in agreement with EU legislation on the general data protection regulation and decisions by the Swedish Ethical Review Authority and Region Skåne, which should be regulated in a material transfer agreement. Enquiries should be directed to.

Code used to create this study will be made available on GitHub upon publication.

## Supplementary Information

**Fig. S1.**
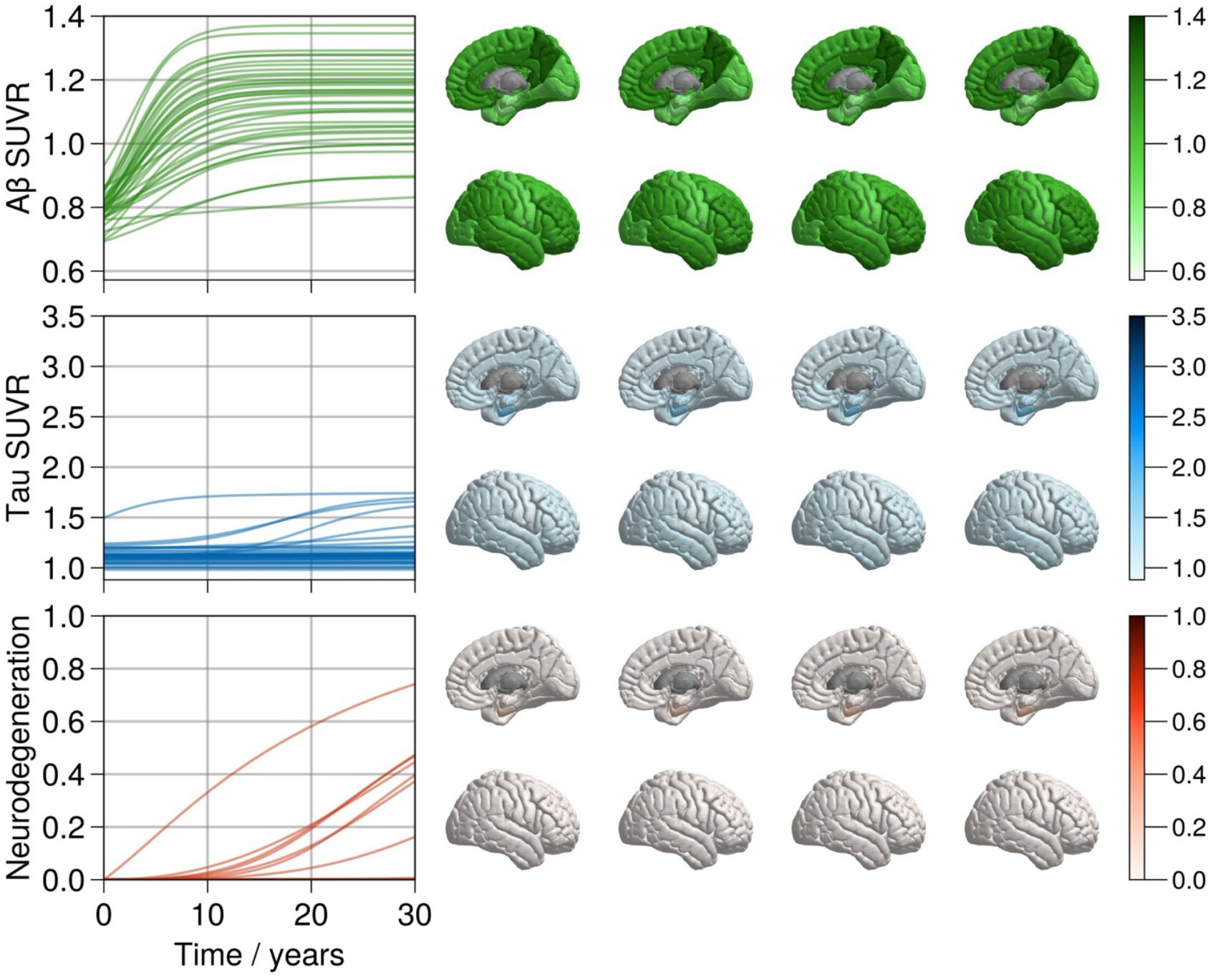
Simulation from the ATN model with no Aβ/tau coupling. Simulation set up is identical to Fig. 1, with the exception that there is no Aβ-tau coupling, β = 0. The trajectories for the right cortex are shown over a 30 year period and cortical renderings on the right cortex at t = {0,10,20,30} years.

**Fig. S2.**
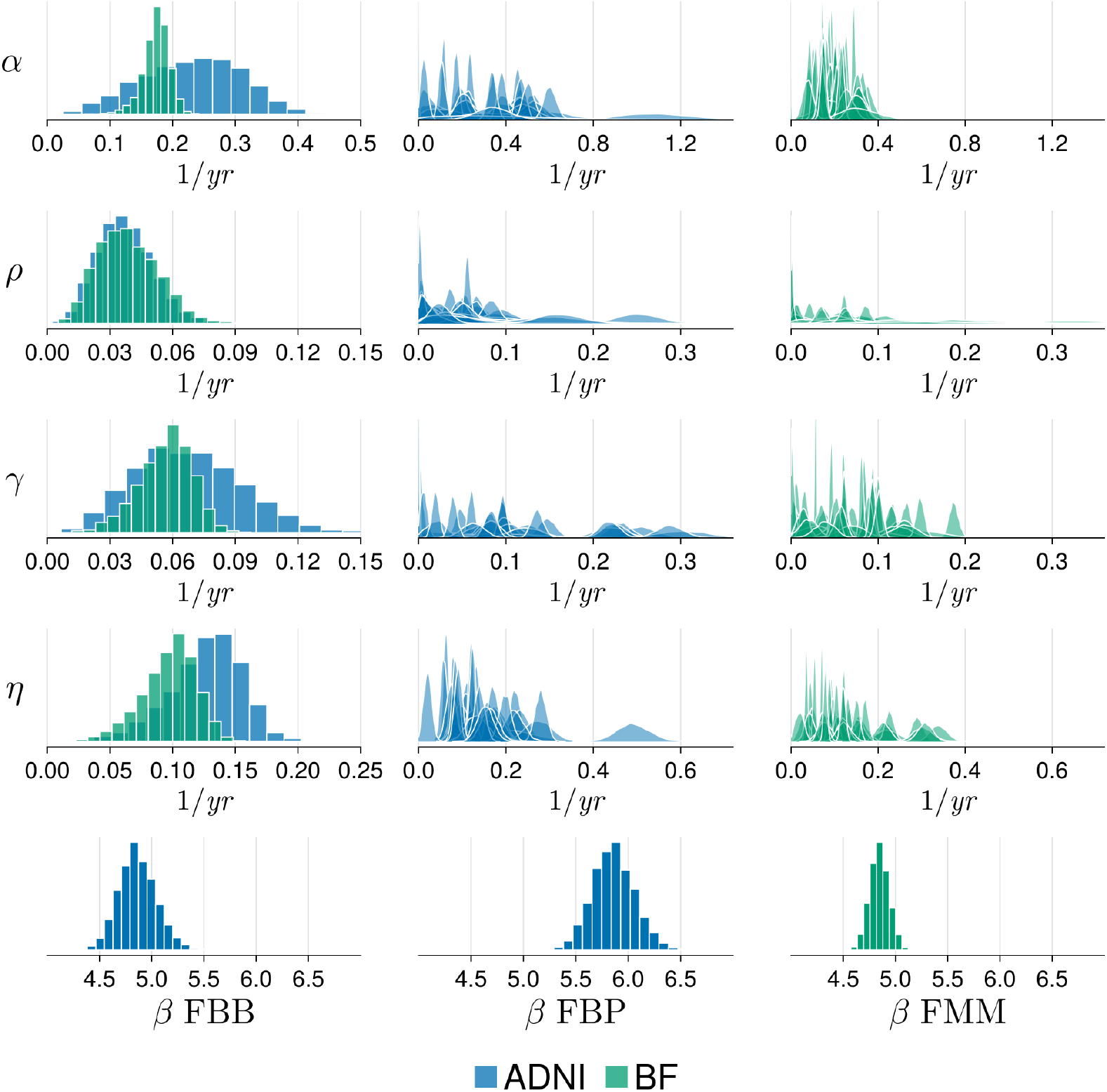
Population-level and individual-level posterior distributions. Posterior distributions from inference on ADNI (blue) and BF2 (green) data for the population-level rate parameters (left panels), and individual-level parameters in ADNI (middle) and BF2 (right). Population-level parameters for the Aβ/tau coupling parameter, β are provided on the bottom row corresponding to florbetaben (FBB), florbetapir (FBP) and flutemetamol(FMM).

**Figure S3.**
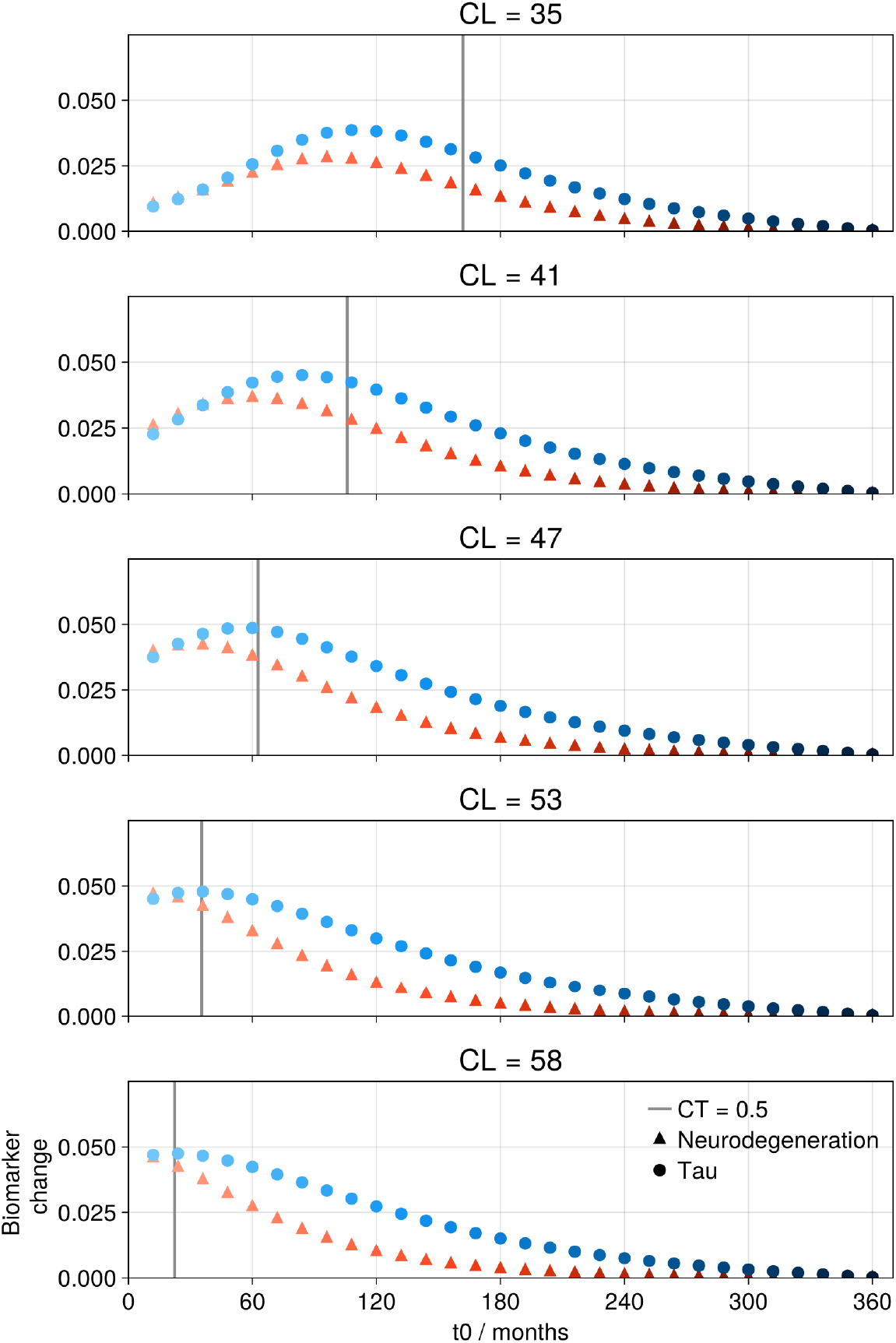
Interactions between endpoint tau and neurodegeneration and colocalisation. Simulated end-point tau and neurodegeneration after a 30-year simulation with parameters in Table 3 and with varying centiloid thresholds defining the cohort from which initial conditions for Aβ and tau. We use thresholds from 60 CL (top panel) to 100 CL (bottom panel) in intervals of 10 CL, the resulting mean CL threshold is given for each cohort, ranging from 35 to 58 CL. For each panel, the colocalisation point, given the initial conditions and parameters, is shown by the solid vertical line. For CL thresholds ≥ 70 CL, the colocalisation point closely coincides with the inflection points in change in end-point neurodegeneration and tau. For CL threshold = 60 CL the colocalisation point occurs after the inflection points. This is likely due to the poor quantification of tau seeds in early disease stages (62).

**Table S1.** Colocalisation order for ADNI. Region and time of Aβ-tau colocalisation for the right and left hemispheres in ANDI, ordered by colocalisation time.

| Right |  | Left |  |
| --- | --- | --- | --- |
| Region | Time / years | Region | Time / years |
| bankssts | 19.18 | inferiortemporal | 16.44 |
| inferiortemporal | 20.67 | fusiform | 19.64 |
| middletemporal | 22.59 | bankssts | 19.72 |
| fusiform | 22.80 | middletemporal | 20.14 |
| parahippocampal | 24.02 | parahippocampal | 20.28 |
| inferiorparietal | 25.07 | entorhinal | 21.03 |
| superiortemporal | 26.66 | inferiorparietal | 22.41 |
| supramarginal | 27.84 | superiortemporal | 24.33 |
| lateraloccipital | 28.67 | Left-Amygdala | 24.61 |
| superiorparietal | 28.97 | temporalpole | 24.92 |
| temporalpole | 29.07 | lateraloccipital | 24.96 |
| Right-Amygdala | 30.22 | supramarginal | 25.66 |
| precuneus | 30.33 | superiorparietal | 26.53 |
| lingual | 31.76 | lingual | 27.30 |
| entorhinal | 32.41 | precuneus | 29.31 |
| isthmuscingulate | 32.69 | insula | 30.46 |
| insula | 33.94 | isthmuscingulate | 31.00 |
| postcentral | 34.97 | rostralmiddlefrontal | 31.92 |
| rostralmiddlefrontal | 35.01 | postcentral | 31.95 |
| lateralorbitofrontal | 35.06 | lateralorbitofrontal | 32.54 |
| parstriangularis | 37.66 | parsopercularis | 32.63 |
| posteriorcingulate | 37.79 | caudalmiddlefrontal | 33.99 |
| cuneus | 38.32 | parstriangularis | 34.53 |
| parsorbitalis | 38.80 | precentral | 35.05 |
| parsopercularis | 38.80 | transversetemporal | 35.83 |
| transversetemporal | 38.88 | cuneus | 36.71 |
| caudalmiddlefrontal | 39.55 | parsorbitalis | 36.81 |
| precentral | 39.90 | superiorfrontal | 36.89 |
| superiorfrontal | 40.04 | posteriorcingulate | 37.67 |
| medialorbitofrontal | 40.96 | pericalcarine | 37.72 |
| pericalcarine | 41.23 | medialorbitofrontal | 40.26 |
| paracentral | 41.34 | rostralanteriorcingulate | 40.81 |
| rostralanteriorcingulate | 41.56 | caudalanteriorcingulate | 42.99 |
| caudalanteriorcingulate | 41.82 | paracentral | 43.24 |
| frontalpole | 48.60 | frontalpole | 47.77 |
| Right-Hippocampus | 55.93 | Left-Hippocampus | 52.97 |

**Table S2.** Colocalisation order for BF2. Region and time of Aβ-tau colocalisation for the right and left hemispheres in BF2, ordered by colocalisation time.

| Right |  | Left |  |
| --- | --- | --- | --- |
| Region | Time / years | Region | Time / years |
| inferiortemporal | 12.61 | inferiortemporal | 12.11 |
| middletemporal | 13.50 | fusiform | 12.90 |
| fusiform | 13.59 | middletemporal | 13.06 |
| bankssts | 14.77 | entorhinal | 13.86 |
| entorhinal | 16.06 | bankssts | 14.21 |
| inferiorparietal | 16.19 | superiortemporal | 15.96 |
| superiortemporal | 16.69 | inferiorparietal | 16.44 |
| lateraloccipital | 18.10 | parahippocampal | 16.85 |
| parahippocampal | 18.22 | Left-Amygdala | 17.73 |
| supramarginal | 18.58 | supramarginal | 17.98 |
| Right-Amygdala | 19.22 | lateraloccipital | 18.17 |
| superiorparietal | 19.34 | temporalpole | 18.52 |
| precuneus | 19.65 | precuneus | 19.00 |
| temporalpole | 19.89 | superiorparietal | 19.71 |
| rostralmiddlefrontal | 21.72 | rostralmiddlefrontal | 20.77 |
| lingual | 22.16 | lingual | 20.80 |
| Right-Hippocampus | 22.44 | isthmuscingulate | 20.81 |
| insula | 22.79 | insula | 21.96 |
| isthmuscingulate | 22.95 | parstriangularis | 22.10 |
| lateralorbitofrontal | 23.43 | parsopercularis | 22.17 |
| postcentral | 23.65 | Left-Hippocampus | 22.26 |
| parstriangularis | 23.95 | caudalmiddlefrontal | 22.82 |
| parsopercularis | 24.29 | postcentral | 22.92 |
| caudalmiddlefrontal | 24.68 | lateralorbitofrontal | 23.19 |
| posteriorcingulate | 24.95 | pericalcarine | 24.02 |
| superiorfrontal | 25.41 | precentral | 24.46 |
| parsorbitalis | 25.49 | superiorfrontal | 24.49 |
| transversetemporal | 25.62 | posteriorcingulate | 24.69 |
| pericalcarine | 25.84 | transversetemporal | 24.83 |
| Precentral | 26.13 | cuneus | 25.83 |
| medialorbitofrontal | 26.92 | parsorbitalis | 25.97 |
| caudalanteriorcingulate | 27.02 | caudalanteriorcingulate | 27.11 |
| cuneus | 27.08 | rostralanteriorcingulate | 27.30 |
| paracentral | 27.11 | paracentral | 28.35 |
| rostralanteriorcingulate | 27.13 | medialorbitofrontal | 28.55 |
| frontalpole | 31.68 | frontalpole | 32.87 |

**Table S3.** Colocalisation Probabilities in ADNI. Colocalisation probability for the right and left hemispheres from ADNI.

| Right |  | Left |  |
| --- | --- | --- | --- |
| Region | Probability | Region | Probability |
| bankssts | 0.852 | inferiortemporal | 0.939 |
| inferiortemporal | 0.144 | parahippocampal | 0.047 |
| inferiorparietal | 0.002 | bankssts | 0.011 |
| parahippocampal | 0.001 | inferiorparietal | 0.004 |

**Table S4.** Colocalisation Probabilities in BF2. Colocalisation probability for the right and left hemispheres from BF2.

| Right |  | Left |  |
| --- | --- | --- | --- |
| Region | Probability | Region | Probability |
| inferiortemporal | 0.855 | inferiortemporal | 0.888 |
| bankssts | 0.094 | bankssts | 0.096 |
| middletemporal | 0.051 | middletemporal | 0.016 |

## Notes

https://adni.loni.usc.edu/

https://biofinder.se/

